# Denoising Single-Cell RNA-Seq Data with a Deep Learning-Embedded Statistical Framework

**DOI:** 10.1101/2025.05.20.655104

**Authors:** Qinhuan Luo, Yongzhen Yu, Tianying Wang

## Abstract

Single-cell RNA sequencing (scRNA-seq) provides extensive opportunities to explore cellular heterogeneity, but is often limited by substantial technical noise and variability. To overcome these issues, we present ZILLNB, a novel computational framework that integrates zero-inflated negative binomial regression with deep generative modeling to simultaneously denoise and impute scRNA-seq data. By integrating a deep generative model, ZILLNB effectively captures latent group structures at both the cellular and gene levels, thereby enhancing the accuracy of recovered gene expression profiles while retaining cell-type-specific expression patterns. Comparative evaluations demonstrate that ZILLNB outperforms existing methods in accurately identifying differentially expressed genes and delineating biologically meaningful cell subpopulations. Moreover, ZILLNB exhibits versatile applicability, including effective correction of batch effects, making it a broadly useful tool for enhancing scRNA-seq data analyses.

## 1 Introduction

Single-cell RNA sequencing (scRNA-seq) technology has significantly advanced our ability to explore diverse cell populations, facilitating the discovery of novel cell types and states [1], elucidating relationships between previously characterized and newly identified cellular phenotypes [2, 3], and improving our understanding of transcriptomic variation in health and disease contexts [4, 5]. However, despite these advantages, scRNA-seq data frequently contain substantial technical noise and variability arising from multiple sources [6, 7, 8]. Specifically, heterogeneity can be categorized into cell-specific, gene-specific, and experiment-specific variations. Cell-specific measurement errors, notably those related to variability in library sizes, pose a significant challenge. These variations primarily stem from differences in sequencing depth and can occur even among cells considered biologically similar [9]. Another prevalent issue is the high frequency of zero counts in scRNA-seq data, often resulting from technical dropout events with distinct patterns across different cell types [10]. Gene-specific errors add further complexity, resulting from interactions between genes or between genes and environmental factors that remain inadequately captured or modeled [11, 12]. Experiment-specific variability, on the other hand, often originates from biases inherent in experimental procedures, such as PCR amplification bias.

Addressing these diverse sources of variability is essential to ensure reliable downstream analyses. Unlike bulk RNA sequencing data, scRNA-seq datasets require specialized computational methods to manage their unique characteristics, particularly the abundance of zeros and associated artifacts. Existing methods generally fall into two main categories: statistical modeling approaches and deep learning-based methods. Statistical approaches, including scImpute [13], VIPER [14], SAVER [15], and ALRA [16], utilize probabilistic frameworks explicitly designed to accommodate the zero-inflation and count distributions inherent to scRNA-seq data. Deep learning-based techniques, such as DCA [17] and DeepImpute [18], exploit neural network architectures, particularly autoencoders, to capture complex, nonlinear relationships among genes. Although these deep learning models provide significant flexibility and scalability, they can suffer from interpretability issues and susceptibility to overfitting, especially when sample sizes are limited.

To leverage the advantages of both statistical modeling and deep learning approaches, we introduce the zero-inflated latent factors learning-based negative binomial (ZILLNB) model. ZILLNB integrates zero-inflated negative binomial (ZINB) regression with deep latent factor models, providing a unified approach for simultaneously addressing various sources of technical variability (Figure 1). By explicitly modeling latent structures at both cell and gene levels, ZILLNB accurately recovers gene expression signals while preserving biologically meaningful variation. Additionally, ZILLNB is designed for broad applicability, effectively handling datasets with or without explicit covariates. We validate ZILLNB’s effectiveness across multiple analytical tasks, including cell type identification, classification of cell subsets, and differential expression analysis. The combined statistical and deep learning approach of ZILLNB thus offers a robust, interpretable, and adaptable solution for the analysis of scRNA-seq data.

**Figure 1.**
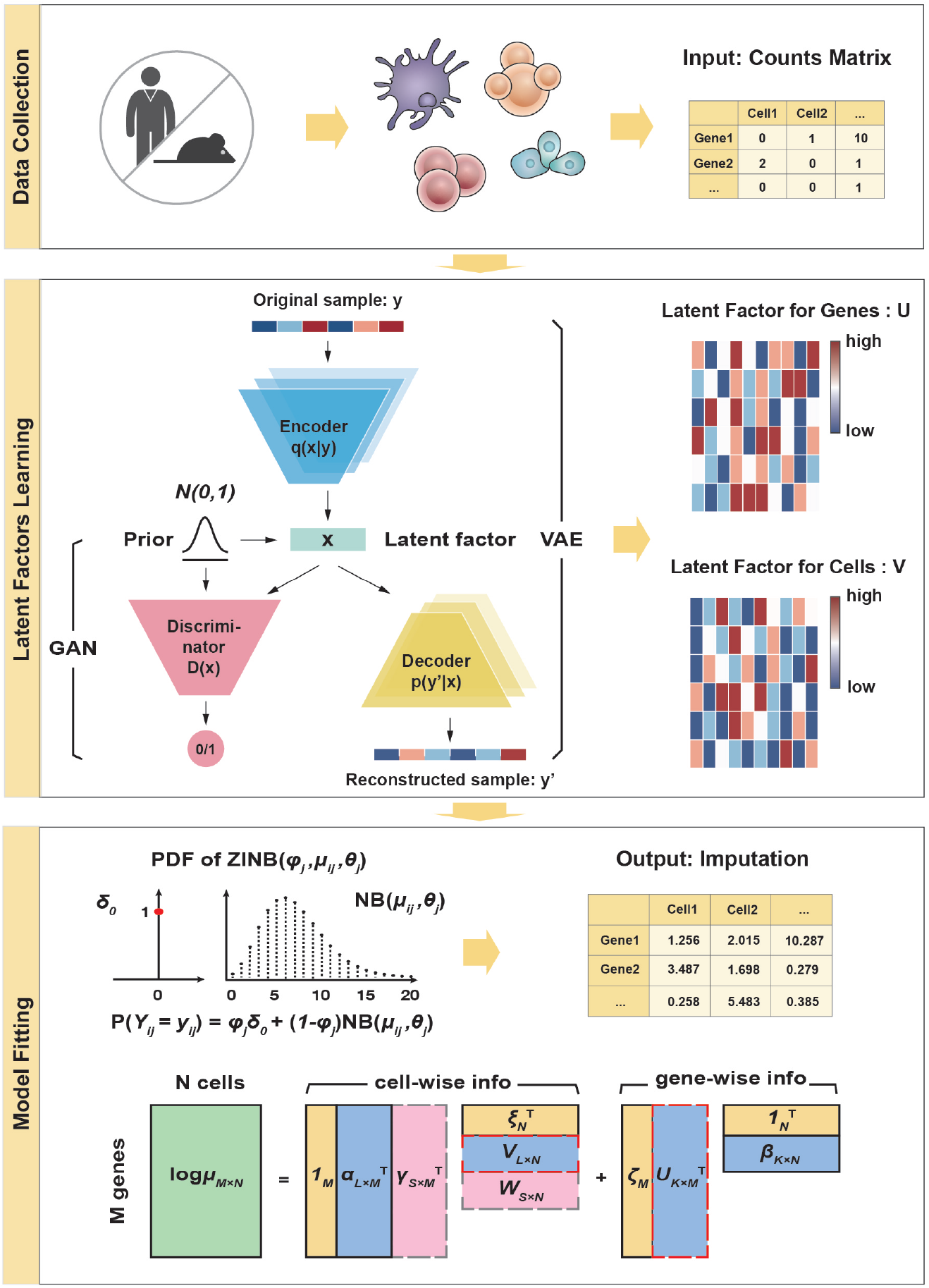
Workflow of the ZILLNB Denoising Method. ZILLNB processes a count matrix using an InfoVAE-GAN model to generate latent factors for cells and genes, which are used to fit a ZINB model. The output is a dense, denoised matrix. The schematic diagram at the bottom shows the decomposition of the link function of the mean value in matrix form. *α*_*L*×*M*_ , *β*_*K*×*N*_ , *γ*_*S*×*M*_ are regression parameters, *ξ*_*N*_ , *ζ*_*M*_ are interception terms, 1_*M*_ , 1_*N*_ are vectors with ones, *U*_*K*×*M*_ , *V*_*L*×*N*_ are latent factors and *W*_*S*×*N*_ is covariates matrix, which is optional. Dimensions *L, K, S* represent the respective sizes of matrices *U, V, W* .

## 2 Materials and methods

The ZILLNB model comprises three main steps (Figure 1). First, an ensemble-based deep generative framework combining InfoVAE and GAN extracts latent features from both cellular and gene-level perspectives. Second, the derived latent factors are utilized to fit a ZINB model, refining both latent representations and regression coefficients through iterative optimization via the Expectation-Maximization (EM) algorithm. This iterative refinement distinctly decomposes variability in the raw data into cell-specific sampling noise and intrinsic gene-specific heterogeneity. Finally, adjusted mean parameters generate a denoised and complete expression matrix.

### 2.1 Latent factors learning

We use the ensemble InfoVAE-GAN model for manifold learning and identifying potentially missing cell-and gene-grouping in scRNA-seq datasets [19, 20]. To reduce overfitting issues common in traditional VAEs, our method integrates InfoVAE with GAN, employing shared network parameters and substituting Kullback-Leibler divergence with Maximum Mean Discrepancy (MMD) [21, 22, 23]:

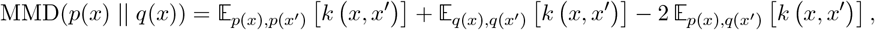

where *p*(*x*), *q*(*x*) denote probability densities, *k*(*x, x*^′^) = exp −|*x* − *x*^′^|^2^*/*2 is the Gaussian kernel, *x*^′^ follows a prior distribution in the latent space, and *x* are latent samples derived from observed data *y*. The InfoVAE-GAN architecture includes three interconnected neural networks: (1) an encoder Enc(*y*) that maps a sample *y* (either a cell or gene vector from the expression matrix *Y* ) to latent space as *x* ∼ Enc(*y*) = *q*(*x*|*y*); (2) a decoder Dec(*x*) reconstructing input samples as *y*^′^ ∼ Dec(*x*) = *p*(*y*^′^|*x*); and (3) a discriminator Dis(*y*), distinguishing real data from generated samples with probabilities defined as *p* = Dis(*y*) ∈ [0, 1]. Adaptive weighting parameters *γ*_1_ and *γ*_2_ balance the reconstruction loss (ℒ_*like*_), prior alignment (ℒ_*prior*_), and generative accuracy (ℒ_*GAN*_ ):

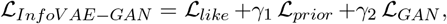

with ℒ_*like*_ = − 𝔼_*q*(*x*|*y*)_[log *p*(*y*|*x*)], ℒ_*prior*_ representing the MMD loss, and ℒ_*GAN*_ = log(Dis(*y*)) + log(1 − Dis(Dec(*x*))) + log(1 − Dis(Dec(Enc(*y*)))). Additional network details are provided in Supplementary Section 1.1.

### 2.2 ZINB fitting

Consider a single-cell RNA sequencing dataset represented by an expression matrix *Y* with dimensions *M* (cells) by *N* (genes). Each element *Y*_*ij*_ represents the observed expression count for gene *j* in cell *i*, modeled using a ZINB distribution. We introduce latent binary variables *Z*_*ij*_ ∼ Bernoulli(*ϕ*_*j*_) indicating whether a zero is generated by a dropout event:

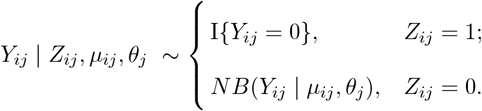

Here, *ϕ*_*j*_ ∈ [0, 1] denotes the gene-specific dropout probability, while *µ*_*ij*_ ∈ ℝ _+_ and *θ*_*j*_ ∈ ℝ _+_ are parameters representing the mean and dispersion of the negative binomial distribution, respectively. The indicator function I(·) equals 1 if its argument is true and 0 otherwise. Following generalized linear modeling, we express the mean parameter *µ*_*ij*_ through a log-link function, incorporating latent cell-and gene-specific structures

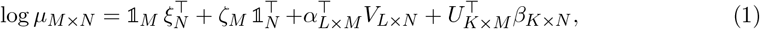

where *U* ∈ ℝ ^*K*×*M*^ and *V* ∈ ℝ ^*L*×*N*^ represent latent factor matrices associated with genes and cells, respectively. Parameters *ξ* ∈ ℝ ^*N*^ and *ζ* ∈ ℝ ^*M*^ account for gene- and cell-specific intercepts, while regression parameters are encapsulated in matrices *α* ∈ ℝ ^*L*×*M*^ and *β* ∈ ℝ ^*K*×*N*^ . Unlike traditional ZINB regression models, which rely on predefined covariates, our approach iteratively updates latent matrices *U* obtained from deep generative models for enhanced fitting (see Figures 4G and S5 for details). In practice, the model converges after a few iterations (Figure S1).

To avoid overfitting, we incorporate regularization terms for the latent factor matrix *U* and intercepts *ξ, ζ* into the model’s objective function, yielding

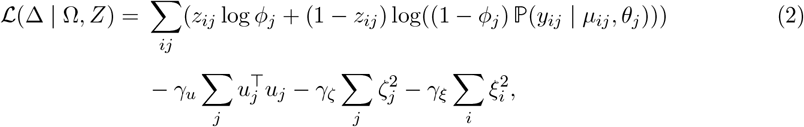

where Ω = {*Y, V* } represents observed data and latent factors, and parameters ∆ include dropout probabilities *ϕ*_*j*_, dispersion parameters *θ*_*j*_, intercepts *ξ*_*i*_, *ζ*_*j*_, regression coefficients *α*_*j*_, *β*_*i*_, and latent gene factors *u*_*j*_, where *i* ∈ {1, · · · , *N* } and *j* ∈ {1, · · · , *M* }. Hyperparameters *γ*_*u*_, *γ*_*ζ*_, *γ*_*ξ*_ control the regularization strength.

We use the EM algorithm, similar to classical ZINB fitting procedures, to iteratively estimate these parameters. Detailed derivations and computations for latent variable updates 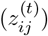 and the EM auxiliary function (*Q*^(*t*)^(∆ | Ω, *Z*)) are provided in Supplementary Section 1.2. Furthermore, external covariates can be included by extending equation (1) with an additional term, 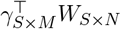 , where *W* ∈ ℝ ^*S*×*M*^ represents external covariate data, *γ* ∈ ℝ ^*S*×*M*^ are the corresponding regression coefficients, and *S* represents the number of covariates. During optimization, since *V* remains fixed, we concatenate *V* and *W* into a combined matrix 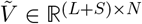 , and similarly merge *α* and *γ* into 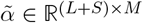 . For notational simplicity, we continue using *V* and *α* to represent these combined matrices throughout the remainder of the manuscript.

### 2.3 Data imputation

Based on latent factors learning and ZINB fitting steps stated above, we introduce the concept of *adjusted mean* as follows: 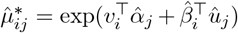, where *v*_*i*_ is the *i*th column of the fixed cell latent factor matrix *V*_*L*×*N*_ , and 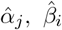, and *û*_*j*_ represent, respectively, the *j*th or *i*th column of the parameters matrices 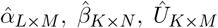 , estimated by minimizing eq (2), with *i* ∈ {1, · · · , *N* } and *j* ∈ {1, · · · , *M* }. By intentionally omitting the intercept terms *ξ*_*i*_ and *ζ*_*j*_, we normalize the original mean parameter *µ*_*ij*_. This normalization allows direct comparability of expression levels across samples while preserving intrinsic biological variability, and is consistent with established denoising practices used in bulk RNA-seq analyses [9].

Additionally, in cases where our goal is to reveal subtle differences between distinct cell types, we further simplify the *adjusted mean* by excluding the 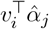 term. The resulting *simplified adjusted mean* is 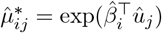, which we denoted as ZILLNB^∗^, a modified variant of the original ZILLNB model.

### 2.4 Dimension reduction and clustering

To effectively visualize high-dimensional raw and imputed data (as shown in Figures 2-6), we adopted the standard workflow from `Seurat` [24] (https://satijalab.org/seurat/articles/pbmc3k_tutorial). Raw data were processed following the default procedures recommended by `Seurat`. For the imputed data, we initially applied column-wise log-normalization, followed by principal component analysis to identify principal components used for dimensionality reduction via Uniform Manifold Approximation and Projection (UMAP). In cases involving large-scale datasets, we additionally employed t-Distributed Stochastic Neighbor Embedding (t-SNE) to enhance visualization quality. Cluster assignments and validations were conducted using hierarchical clustering with average linkage. Notably, clustering analyses performed on the estimated latent matrix *Û* followed the identical approach used for the raw data.

**Figure 2.**
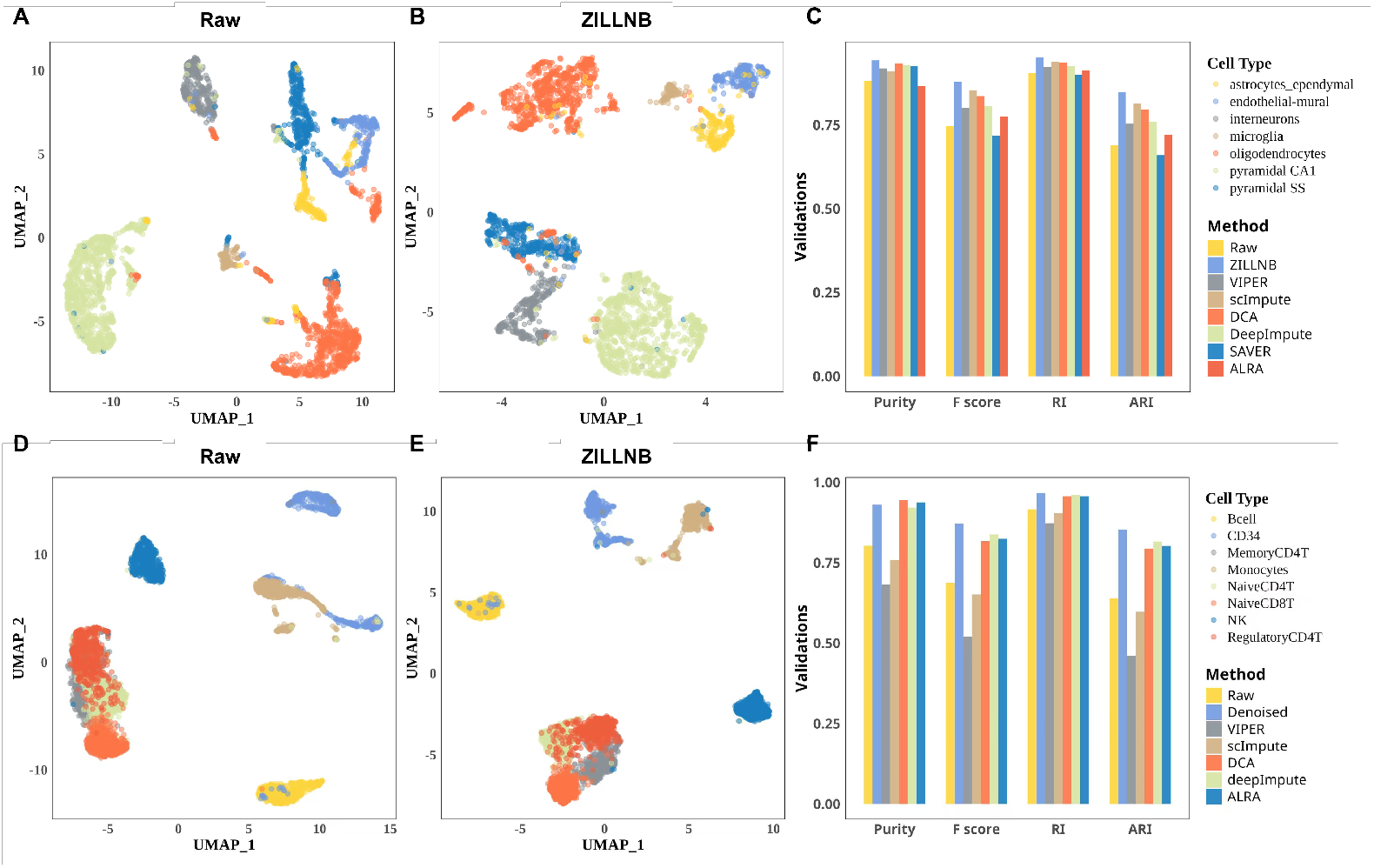
ZILLNB improves cell classification accuracy and compactness in mouse brain cortex and subsampled human PBMC scRNA-seq datasets. (A, B, D, E) UMAP plots of the mouse brain and subsampled human PBMC scRNA-seq data using raw and ZILLNB-imputed data, with points labeled by true cell types. (C, F) Four external clustering validations (Purity, F score, RI, and ARI) demonstrate the performance and robustness of ZILLNB compared to five widely used imputation methods, evaluated on the mouse brain dataset and human PBMC dataset.

To assess clustering accuracy, we calculated four metrics against true cell labels provided in each original dataset: Purity, F score, Rand index (RI), and adjusted Rand index (ARI) [25]. Each metric ranges from 0 to 1, with values closer to 1 indicating superior clustering performance. Detailed definitions and formulas for these metrics are available in Supplementary Section 1.3.

### 2.5 Random matrix comparison

To quantitatively evaluate the learned latent factor matrix *Û*_*K*×*M*_ against random Gaussian noise, we employ spectral methods derived from random matrix theory [26]. Consider a matrix *X* ∈ ℝ ^*D*×*N*^ whose entries are independently drawn from a Gaussian distribution *N* (0, *σ*^2^). We examine its sample covariance matrix *W* = *XX*^⊤^*/N* through three eigenvalue-based metrics: (1) empirical spectral density *µ*_*W*_ , which asymptotically follows the Marchenko-Pastur distribution; (2) empirical normalized level-spacing density *µ*_*G*_, approaching the Wigner Surmise distribution; and (3) the largest eigenvalue (spectral radius) *λ*(*D*), which converges to the Tracy-Widom distribution. Detailed derivations and asymptotic distributions for these statistics are provided in Supplementary Section 1.4. For an accurate comparison, we standardized *Û*_*K*×*M*_ by centering and scaling each column to zero mean and unit variance. Subsequently, we derived its corresponding covariance matrix *W*_*U*_ and calculated the aforementioned spectral statistics. The goodness-of-fit between empirical and theoretical distributions was assessed using Kolmogorov-Smirnov and Anderson-Darling tests. The largest eigenvalue comparison utilized the `RMTstat` package in `R` [27], which directly computes *p*-values based on the Tracy-Widom distribution.

### 2.6 Gene sets evaluation via enrichment analysis and AUCell scores

We assessed gene set relevance through enrichment analysis and gene set activity measurements using the `AUCell` package [28] in `R`. Gene Ontology (GO) and Kyoto Encyclopedia of Genes and Genomes (KEGG) pathway analyses were conducted using the `clusterProfiler` package [29], with the top 10 pathways visualized (Figure S4). To further evaluate biological significance, we calculated AUCell scores to quantify gene cluster activities, subsequently averaging these scores by cell type within each cluster. Results were presented using heatmaps (Figures 4F and S3).

**Figure 3.**
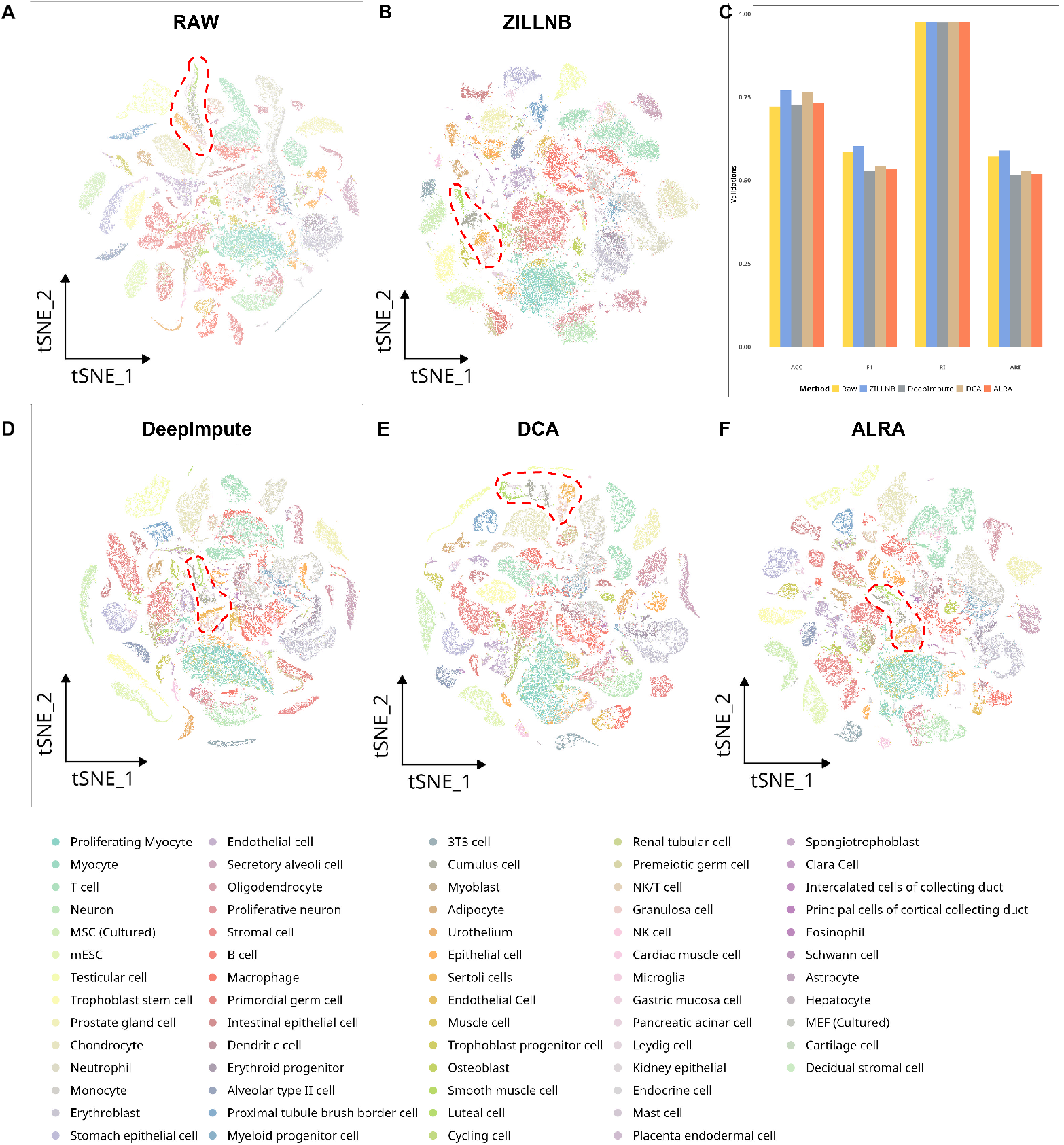
ZILLNB improves resolution and clarity in cell classification within the MCA scRNA-seq dataset. (A, B, D, E, F) tSNE plots displaying cell-type annotations of the mouse cell atlas dataset, before and after denoising using various imputation methods: (A) raw data, (B) ZILLNB, (D) DeepImpute, (E) DCA, and (F) ALRA. (C) Four external clustering validations (Purity, F score, RI, and ARI) confirm ZILLNB’s superior performance compared to other imputation methods.

**Figure 4.**
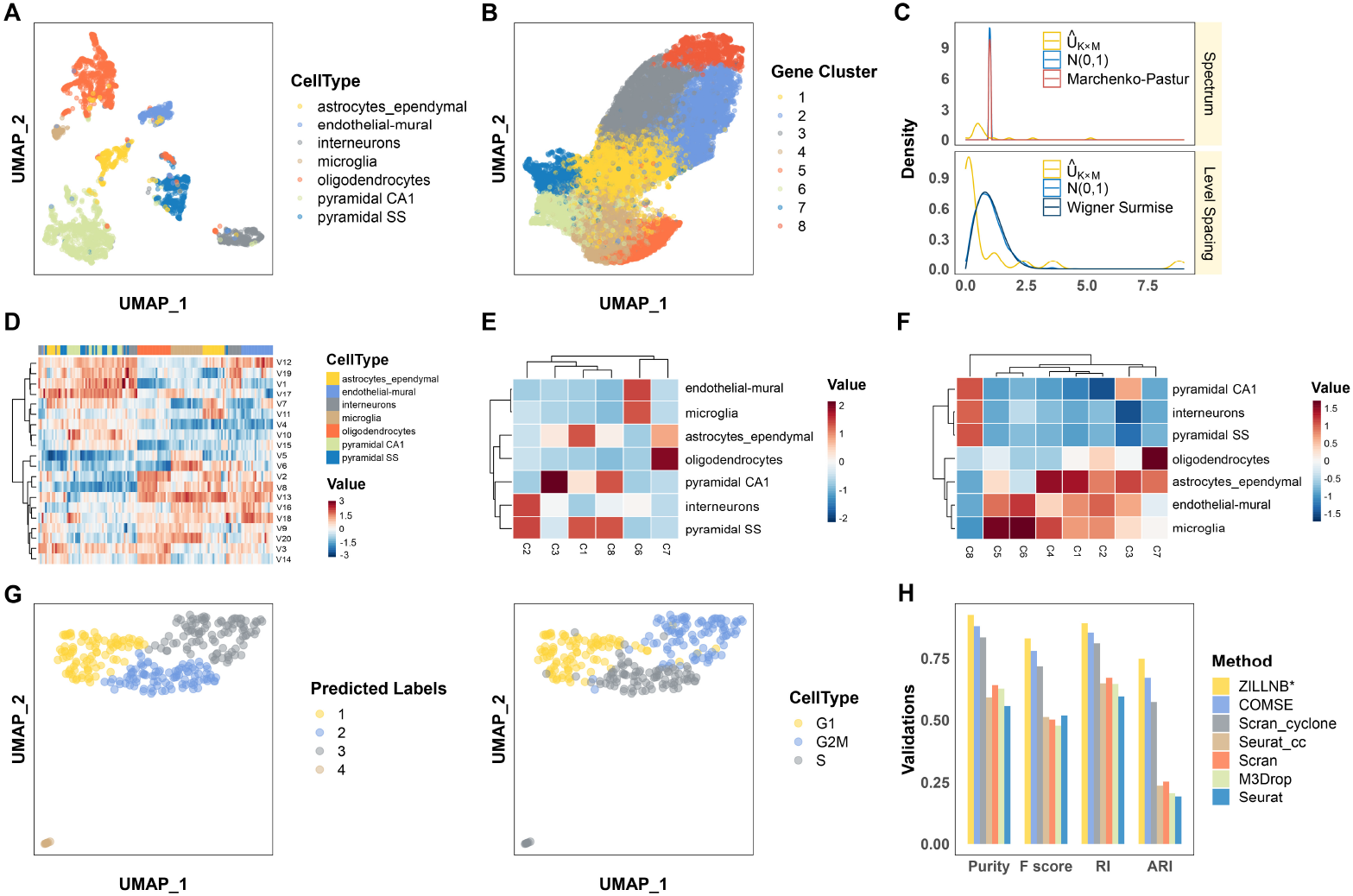
Latent factor learning preserves cell-wise and gene-wise information in the mouse brain dataset [33]. (A, B) UMAP projections of cell latent factors *V* and gene latent factors *Û*, annotated by cell types and Seurat-predicted clusters. (C) Comparison of learned gene latent factor matrix *Û* with a standard Gaussian matrix using Wishart ensemble. The upper panel shows empirical spectral distributions versus the theoretical Marchenko-Pastur law (red curve); the lower panel illustrates the level-spacing distributions alongside the Wigner Surmise law (blue curve). (D) Heatmap of the top 20 marker genes per cell type based on *Û*, with genes as columns and latent variables (*K* = 20) as rows. (E) Heatmap illustrating the overlap between the top 20 marker genes and Seurat-predicted gene clusters. (F) AUCell scores for the predicted gene clusters across various cell types, reflecting activity levels within the original dataset. (G) UMAP projection of the cell cycle dataset [36] based on 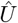, annotated by predicted and true cell types. (H) Clustering performance assessment using Purity, F score, RI, and ARI metrics of ZILLNB^∗^, compared with COMSE [37], Scran cyclone [38], Seurat cell-cycle scoring [39] and regression, Scran [40], M3Drop [41], and Seurat method applied to the cell cycle dataset.

### 2.7 Differentially expressed genes (DEGs) selection and simulation design

ZILLNB accommodates datasets enriched with metadata (e.g., treatment vs. control, disease vs. normal) by integrating one-hot encoded covariates into matrix *V* . To simulate realistic experimental scenarios, we generated subsets from the Mixture 1 breast cell line dataset [30], selecting cells from one tumor type (BT474 or T47D) and one non-tumor type (Jurkat or Thp1) to establish distinct treatment and control groups. These subsets formed the basis for creating 100 bootstrapped datasets utilized in differential expression analyses. For selecting DEGs, we applied established bulk RNA-seq methodologies, specifically DESeq2, edgeR, and limma, employing standard criteria for false discovery rate (FDR) and log2 fold change (logFC). Both log-normalized raw data (processed using `Seurat`) and ZILLNB-denoised data were subjected to statistical testing (t-tests and Wilcoxon tests), based on tumor and non-tumor classifications. FDR correction was performed using the Benjamini-Hochberg procedure [31], and logFC values were computed for each comparison.

To simplify threshold selection for identifying DEGs, we combined FDR and logFC values into a unified ranking metric, *f* , representing the likelihood of a gene being a genuine DEG. Specifically, we used quantiles derived from ascending FDR values and descending absolute logFC as boundaries, constructing a quantile-based grid with increments of 10^−3^. The diagonal line of this grid defined the nested threshold boundary, where genes positioned within regions

of lower FDR and higher absolute logFC were classified as predicted DEGs (pDEGs). For bulk RNA-seq data, we enhanced reliability by intersecting DEGs derived independently from DESeq2, edgeR, and limma, producing stable and robust true DEGs (tDEGs) [32]. Genes in the difference set of adjacent squares were ranked by counting the total number of genes within the larger quantile squares, normalized by the total gene count, resulting in the score *f* .

Following the generation of tDEGs and pDEGs sets, we assessed performance using conventional metrics: FDR, True Positive Rate (TPR), Accuracy (ACC), and ARI, as detailed in Supplementary Materials. Additionally, by varying the number of genes selected as pDEGs, we constructed Receiver Operating Characteristic (ROC) curves and computed Area Under the Curve (AUC) values, with tDEGs serving as a fixed reference set. In practical terms, we considered the top 20% of bulk RNA-seq genes (*f* = 0.2, approximately 3,000 genes) as the tDEGs for sub-figures A, B, D, and E in Figures 5 and S6, replicating realistic scenarios. Robustness was further evaluated by systematically adjusting *f* values from 0 to 0.5, measuring corresponding AUC values, Pearson correlation, Wasserstein-2 distance, and cross-entropy, as described comprehensively in the Supplementary section 1.3.

**Figure 5.**
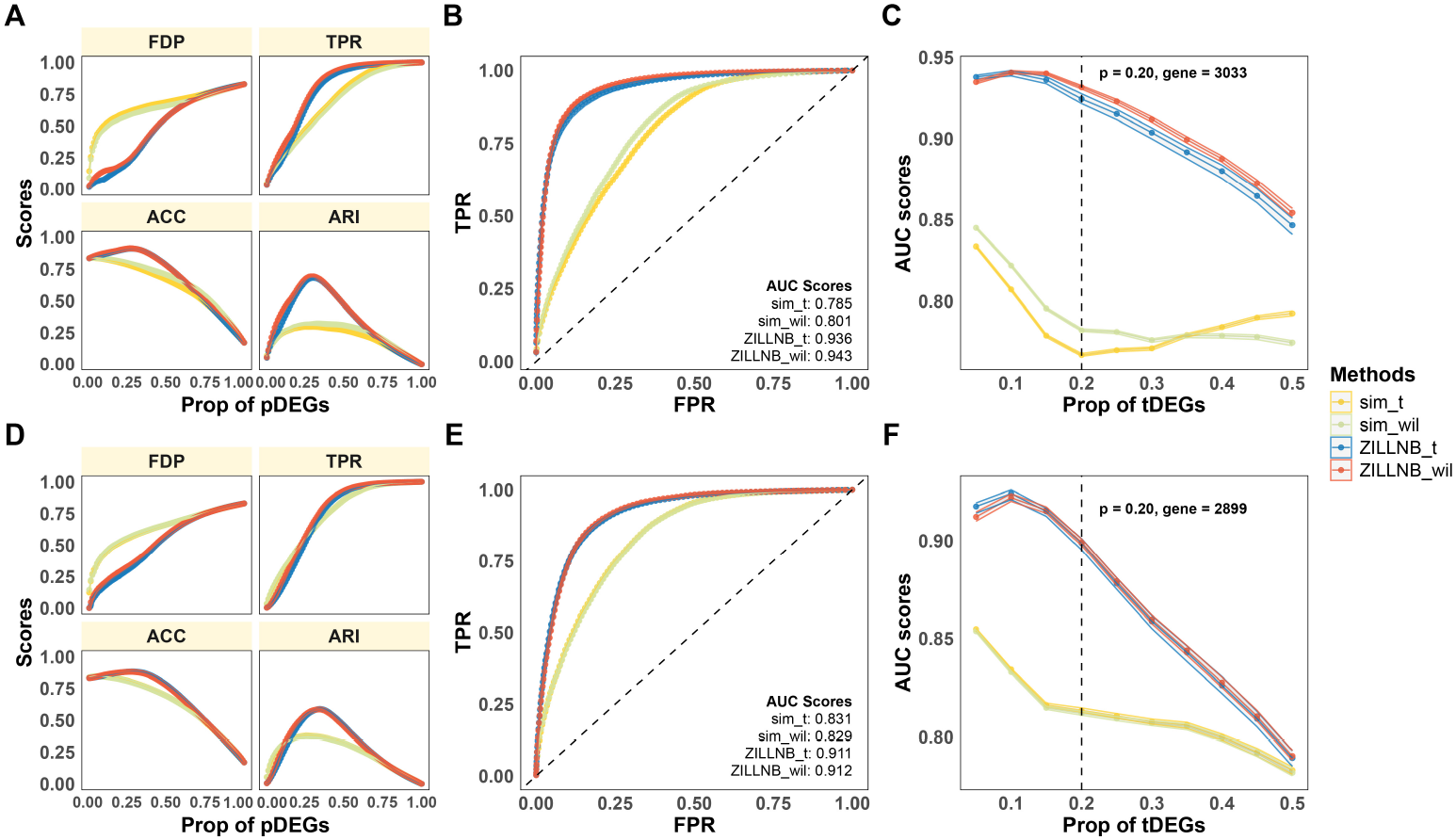
ZILLNB robustly identifies DEGs in bootstrap experiments using breast cancer cell line dataset [30]. Rows correspond to comparisons between cell lines: the first row compares BT474 versus Jurkat, and the second row compares BT474 versus Thp1. (A, D) Evaluation metrics: FDP, TPR, ACC, and ARI. (B, E) ROC curves generated using fixed proportions of tDEGs identified from bulk RNA-seq data, compared against varying proportions of pDEGs. (C, F) AUC scores for varying proportions of tDEGs from 0 to 0.5, with dashed lines marking the scenario of 20% tDEGs (approximately 3,000 genes). Methods sim t/sim wil represent the log-normalized Seurat method combined with t-tests/Wilcoxon tests, while ZILLNB t/ZILLNB wil uses the denoised ZILLNB matrix combined with the t/Wilcoxon tests.

## 3 Results

### 3.1 Enhanced cell classification after denoising

To evaluate the effectiveness of the ZILLNB framework, we compared its performance against several widely utilized imputation methods, including SAVER [15], scImpute [13], VIPER [14], DeepImpute [18], and DCA [17]. Specifically, we assessed the accuracy of each method in recovering missing transcript counts and preserving biologically relevant signals critical for accurate cell-type classification.

Our evaluations involved two scRNA-seq datasets with verified cell-type annotations: (1) Mouse Brain Dataset: consisting of 3,005 single cells obtained from the cerebral cortex of 33 male and 34 female mice (GSE60361) [33]; and (2) Human PBMC Dataset: comprising eight distinct immune cell types from purified human peripheral blood mononuclear cells (PBMCs), subsampled to 500 cells per cell type [34]. We compared predicted labels derived from each imputation method against true cell-type labels, employing four clustering performance metrics: Purity, F score, RI, and ARI (see the Methods and the Supplementary section 1.3).

The ZILLNB framework consistently outperformed standard imputation methods in clustering and cell type classification across all datasets and metrics, demonstrating improved Purity, F score, RI, and ARI (Figures 2C, F). Notably, ZILLNB demonstrated a significant improvement in ARI, highlighting its robustness in accurately identifying cell-type groupings. Methods such as VIPER and scImpute did not outperform when using the raw data directly, likely due to challenges in distinguishing subtle subgroup structures within highly similar cell populations, such as the T cells in the PBMC dataset (Figure 2F).

To further illustrate the capability of ZILLNB in analyzing large-scale scRNA-seq datasets, we employed the mouse cell atlas (MCA) dataset [35], which includes samples derived from more than 50 mouse tissues and cultures. Compared with DeepImpute, DCA, and ALRA [16], ZILLNB showed a t-SNE representation with superior resolution of subcell types (Figure 3). Specifically, gonadal-related cell subtypes (including luteal cells, cumulus cells, granulosa cells, Leydig cells, and Sertoli cells) exhibited clearer and more distinct boundaries after ZILLNB denoising, as indicated by the red dashed circles in Figure 3B. Moreover, cluster evaluation metrics further confirmed the improved performance of ZILLNB (Figure 3C).

By showing the accurate cell clustering results, ZILLNB effectively preserves latent biological characteristics while capturing meaningful underlying signals.

### 3.2 Latent factors preserve cell- and gene-wise information

To assess the biological and statistical significance of the latent factor matrices *U* and *V* produced by ZILLNB, we evaluated their ability to retain meaningful information from multiple perspectives. The matrix *V* effectively captured the underlying structure of cell populations, as evidenced by Figures 4A. Additionally, we verified the capability of matrix *V* to accurately preserve biological signals despite significant batch effects present in the raw data. By integrating batch information encoded as a one-hot matrix into the *V* matrix training process using pancreatic scRNA-seq datasets generated from diverse sequencing technologies [42] (Figure S2A), *V* effectively mitigated sequencing biases, achieving superior clustering accuracy compared to methods like Seurat-CCA [24] and Harmony [43] (Figures S2B, C, D, E).

For the gene-embedding latent factor matrix *Û*_*K*×*M*_ , estimated from eq (2), we validated its capacity to capture biologically relevant gene structures using the mouse brain dataset [33], using both statistical and biological analyses. We compared *Û* against a random Gaussian matrix *X* of equivalent dimensions, sampled from *N* (0, 1), the prior distribution used during training. Spectral density analysis, normalized level-spacing distributions, and largest eigenvalue assessments revealed pronounced distinctions between *Û* and the random matrix *X* (Figure 4C). Goodness-of-fit tests (Kolmogorov-Smirnov and Anderson-Darling tests, *p <* 10^−14^) and the largest eigenvalue test (*p* = 0) strongly indicated that *Û* encodes structured latent signals rather than random noise.

The biological relevance of *Û* was further supported by clustering and enrichment analyses. Using Seurat’s clustering pipeline, we identified eight gene clusters and examined their biological significance (Figure 4D). The heatmap depicting the top 20 marker genes per cell type confirmed that *Û* retained critical marker gene information, with clearly identifiable clusters for microglia, oligodendrocytes, and pyramidal cells. Additional analyses indicated strong enrichment of marker genes within specific gene clusters (Figures 4E, S3). GO and KEGG enrichment analyses validated cell-type-specific functions, highlighting immune-related processes in microglia and differentiation-related functions in oligodendrocytes (Figure S4). AUCell analysis further confirmed high activity of gene clusters 6, 7, and 8 within microglia, oligodendrocytes, and neurons, respectively (Figure 4F).

We also validated ZILLNB’s effectiveness using a homogeneous mouse embryonic stem cell (mESC) dataset [36], where nuanced gene-level variations define distinct cell cycle phases (G1, G2M, S). Utilizing the 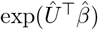 representation, ZILLNB clearly differentiated these cell cycle phases (Figure 4G and S5), demonstrating superior clustering performance compared to methods such as COMSE [37], Scran Cyclone [38], and Seurat cell-cycle scoring [39] (Figure 4H).

### 3.3 ZILLNB enhances DEGs selection

To illustrate the utility of ZILLNB in downstream analyses, we evaluated its performance in detecting DEGs, a critical task in scRNA-seq data analysis. Since ground-truth DEG labels are typically unavailable for scRNA-seq datasets, we employed complementary scRNA-seq and bulk RNA-seq datasets. The latter provided benchmark DEG labels for robust validation. Performance stability was rigorously assessed through 100 bootstrap iterations for each experimental dataset (see Methods 2.7 for details).

We analyzed a breast cancer cell line dataset containing both scRNA-seq and bulk RNA-seq data for each tumor and non-tumor cell line [30]. Semi-synthetic datasets were created by combining one tumor (BT474 or T47D) with one non-tumor cell line (Jurkat or Thp1) to simulate treatment and control groups. DEG selection was performed using established criteria involving FDR and logFC. tDEGs were defined by intersecting results obtained independently from DESeq2, edgeR, and limma analyses on bulk RNA-seq data [44, 9, 45].

Using the top 20% (∼3,000 genes) of bulk RNA-seq data as benchmark tDEGs, ZILLNB consistently surpassed the baseline log-normalization approach across multiple evaluation metrics. ZILLNB achieved significantly reduced False Discovery Proportion (FDP) and enhanced TPR, clearly demonstrating its superior recall capabilities (Figure 5A, S6). Furthermore, ZILLNB yielded superior ROC curves and higher AUC scores compared to the naive t-test and Wilcoxon test (Figure 5B). When the proportion of tDEGs was systematically varied from 0% to 50% ( 7,500 genes), ZILLNB maintained consistent and substantial improvements over the baseline method across all scenarios (Figure 5C, F, S6). Additional quantitative comparisons using Pearson correlation, Wasserstein-2 distance, and cross-entropy further supported ZILLNB’s superior performance in capturing genuine biological signals from scRNA-seq data (Figure S7). These findings were consistently replicated across other cell-line comparisons (BT474-Jurkat and BT474-Thp1), underscoring the robustness and reliability of ZILLNB (Figure 5D-F, S6, S7).

Overall, ZILLNB demonstrated robust advantages over traditional log-normalization methods, achieving improved FDP control, higher TPR, and enhanced preservation of intrinsic data structure. Its consistently strong performance across diverse scenarios and perturbations underscores ZILLNB’s effectiveness and reliability for DEG identification in scRNA-seq analyses.

### 3.4 ZILLNB facilitates identification of fibroblast subsets

We next demonstrate ZILLNB’s capability in identifying specific fibroblast subsets undergoing fibroblast-to-myofibroblast transition (FMT), a crucial event in idiopathic Pulmonary Fibrosis (IPF). Utilizing ZILLNB-denoised scRNA-seq data, we identified a distinct cluster exhibiting robust myofibroblast-associated characteristics, validating its biological significance through marker gene expression, pathway enrichment, and gene set activity analyses.

IPF, the most common type of pulmonary fibrosis, is characterized by chronic progressive lung fibrosis [46]. The FMT drives myofibroblast accumulation, and inhibiting this transition can significantly mitigate or halt disease progression [47, 48]. Given the critical role of myofibroblasts in IPF pathogenesis, accurately identifying fibroblast subsets involved in FMT is essential for advancing therapeutic strategies.

To this end, we analyzed publicly available scRNA-seq datasets of IPF, focusing specifically on fibroblast populations. Using ZILLNB, we identified a clearly defined fibroblast subset (cluster 3) characterized by elevated expression of myofibroblast markers, including MYO1E, H19, COL3A1, and COL1A1 (Figure 6A). We further evaluated these cells by generating a curated myofibroblast-related gene list from the GeneCards database [49], applying a relevance score threshold of ⩾ 1, resulting in a set of 517 genes. AUCell scoring analysis confirmed enriched expression of these myofibroblast-associated genes within cluster 3 (ZILLNB) (Figure 6A).

**Figure 6.**
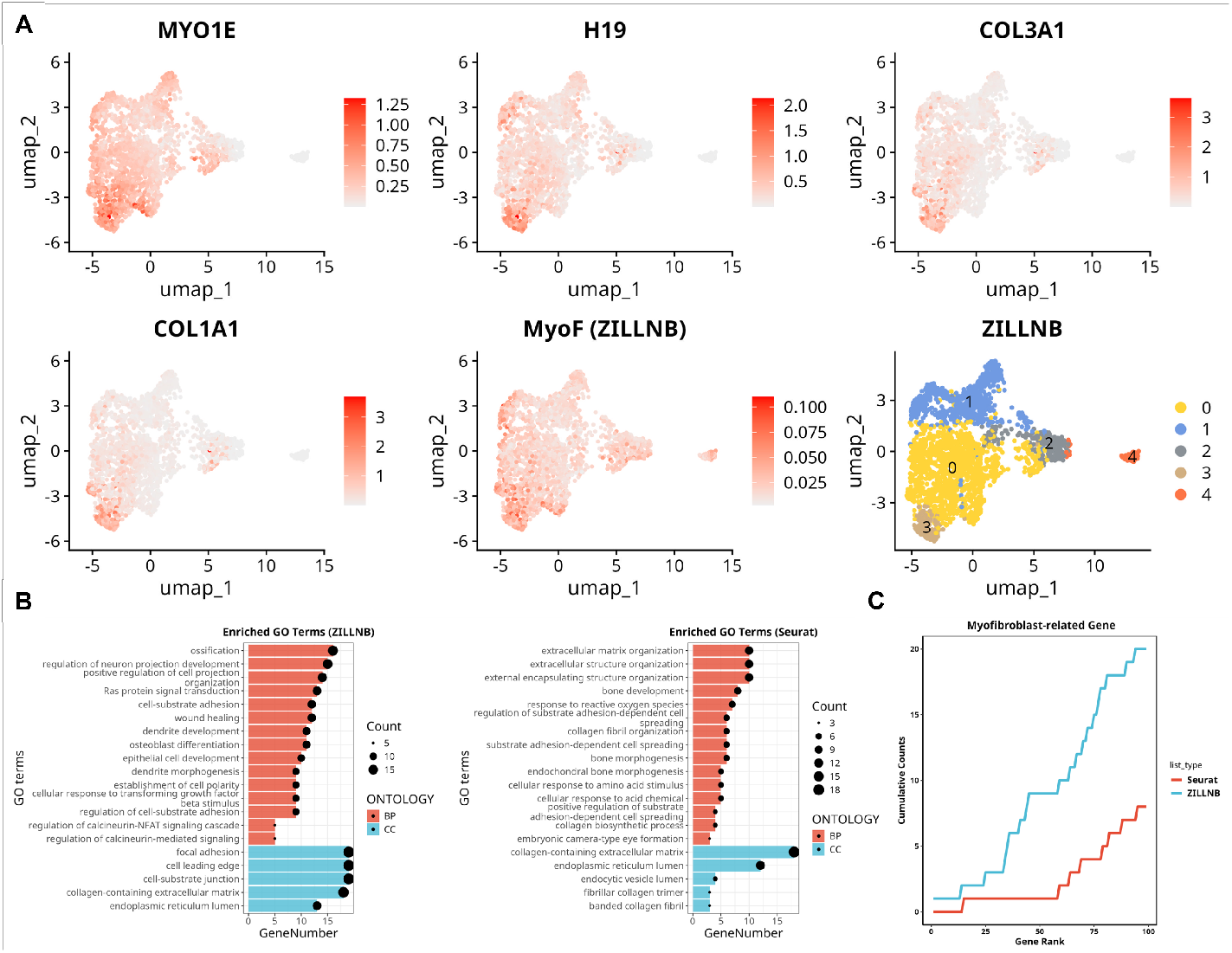
Identification of fibroblast subsets undergoing FMT. (A) UMAP plots of re-clustered fibroblasts from 3 patients with IPF, 3 patients with chronic obstructive pulmonary disease (COPD), and 3 healthy controls, displaying expression levels of myofibroblast markers, AUCell scores for myofibroblast-associated gene sets, and Seurat-predicted labels derived from ZILLNB-denoised data. (B) GO analysis of the top 200 DEGs in cluster 3 identified by ZILLNB compared to the Seurat pipeline. The bar plot illustrates the number of DEGs associated with each GO term across Biological Process (BP), Cellular Component (CC), and Molecular Function (MF). (C) Cumulative curve showing the top 200 DEGs in cluster 3 (ZILLNB) and cluster 4 (Seurat) that overlap with myofibroblast-related gene sets.

In contrast, a parallel analysis using the raw data and the standard Seurat pipeline identified another cluster (cluster 4) with similar marker gene expression. However, this cluster exhibited less coherent signals, and AUCell scoring failed to distinctly identify a myofibroblast-enriched subset (Figure S8A). These observations underscore ZILLNB’s superior precision in accurately delineating biologically relevant cell subsets.

To further establish biological validity, we conducted GO enrichment analysis of the top 200 DEGs from cluster 3 identified by ZILLNB. Results revealed significant enrichment in wound healing processes, a characteristic functional signature of myofibroblasts (Figure 6B). Additionally, cumulative curve analyses showed that ZILLNB-derived DEGs encompassed a higher proportion of myofibroblast-related genes compared with those identified via the raw Seurat analysis, further highlighting the biological significance of the identified subset (Figure 6C).

Thus, these results affirm that ZILLNB provides clearer and more biologically accurate identification of fibroblast subsets undergoing FMT. By effectively denoising scRNA-seq data, ZILLNB emerges as a robust analytical tool with broad applicability for investigating complex biological phenomena such as IPF and similar multifactorial diseases.

## 4 Discussion

ZILLNB introduces an innovative deep generative modeling framework integrated with classical two-part modeling, effectively transforming the traditionally unsupervised task of modeling scRNA-seq data into a supervised learning context. By dynamically capturing complex multi-variate structures at both gene and cell levels, ZILLNB merges the strengths of deep learning’s powerful representational capabilities with the interpretability and robustness inherent to ZINB models.

Previous ZINB-based methods have been extensively utilized for scRNA-seq and microbiome analyses. However, these approaches typically depend heavily on user-defined covariates or limit covariate dimensionality, restricting the complexity of captured group structures [50, 51, 52]. In contrast, ZILLNB employs neural network architectures to autonomously learn intricate gene-wise and cell-wise structures without predetermined covariate constraints. Critically, ZILLNB does not merely feed neural network-derived covariates into a static ZINB model; rather, the gene latent matrix *U* is dynamically updated throughout training, whereas the cell latent matrix *V* remains fixed after adequately capturing cell-wise structure (Figure 4A, S2).

This integrated training strategy reduces computational complexity and significantly enhances representational capacity compared to traditional ZINB implementations. Thus, by incorporating deep learning principles, ZILLNB effectively expands the applicability of classical statistical models, particularly when crucial biological information is indirectly observed or latent.

Although initially designed to address heterogeneity in scRNA-seq data, ZILLNB demonstrates considerable adaptability across a broad spectrum of analytical applications. For instance, it can readily integrate user-specified covariates, making it highly effective in tasks such as batch-effect correction and cohort bias adjustment. Users can also define particular gene and cell groups to specifically target variance sources or amplify relevant expression signals. As demonstrated in Figure 5, defining tumor and normal cell groups significantly enhances robust DEG identification. Similarly, ZILLNB’s flexibility enables the modeling of diverse structures, including cell types, experimental conditions, and biological perturbations, to enhance downstream analyses such as precise cell subset characterization (Figure 6).

ZILLNB represents a versatile analytical framework for interpreting complex scRNA-seq data, offering extensive potential applications. Future extensions and adaptations could further broaden its utility. Promising directions include applying ZILLNB to pseudo-time trajectory analyses [2], enabling detailed exploration of dynamic cellular processes across developmental timelines by integrating latent-space similarity measures. Additionally, extending ZILLNB for gene clustering and regulatory network inference [53] could yield valuable insights into gene functionality and interactivity. Furthermore, adapting ZILLNB to bulk RNA-seq or scATAC-seq data would facilitate critical tasks such as data denoising and differential feature identification, underscoring its versatility in analyzing diverse genomic data types.

## 5 Acknowledgements

We would like to thank Dr. Xun Lan from Tsinghua University for generously providing computational resources.

## Authors’ contributions

T.W., Q.L., and Y.Y. conceptualized and designed the model and computational framework. Q.L. and Y.Y. developed the algorithm, implemented the model, and conducted performance evaluations on real datasets. T.W. led the manuscript preparation. All authors provided substantial critical feedback, shaping the research, analysis, and manuscript preparation. T.W. supervised the overall project. All authors reviewed and approved the final manuscript.

## 6 Supplementary data

Supplementary data are available at NAR Genomics & Bioinformatics online.

## 8 Data and code availability

All datasets utilized in this analysis are publicly available. The mouse brain scRNA-seq dataset comprising 3,005 cells from the mouse cortex and hippocampus can be accessed through the Gene Expression Omnibus (GEO) SuperSeries GSE60361 [33]. The human PBMC dataset was subsampled to 4,000 cells from an original 68k PBMC dataset available in the Short Read Archive under accession SRP073767, encompassing proportionally subsampled cells from eight distinct immune cell types [34]. The expression matrix for the MCA dataset is available at https://ndownloader.figshare.com/files/10351110?private\_link=865e694ad06d5857db4b, with annotations accessible via https://ndownloader.figshare.com/files/10760158?private\_link=865e694ad06d5857db4b. The human pancreatic scRNA-seq datasets generated using various sequencing methods [42] can be accessed under accession numbers GSE81076, GSE85241, GSE86469, GSE84133, GSE81608, and E-MTAB-5061 (ArrayExpress). The mESC scRNA-seq datasets, generated using full-length and UMI-based techniques, are available under accession numbers E-MTAB-2805 [36] and GSE54695 [54], respectively. The bulk RNA-seq

dataset for mESCs is accessible under GSE78140 [55]. The breast cancer cell line dataset, comprising both bulk RNA-seq and scRNA-seq data, is publicly accessible under accession GSE220608 [30]. The source code for ZILLNB, implemented in R and Python, is available on FigShare (https://doi.org/10.6084/m9.figshare.28978697, https://doi.org/10.6084/m9.figshare.28978151, and https://doi.org/10.6084/m9.figshare.28978085).

## References

[1] Alev Baysoy, Zhiliang Bai, Rahul Satija, and Rong Fan. The technological landscape and applications of single-cell multi-omics. Nature Reviews Molecular Cell Biology, 10 2023.

[2] Cole Trapnell. Defining cell types and states with single-cell genomics. Genome Research, 25:1491–1498, 10 2015.

[3] Evan Z. Macosko, Anindita Basu, Rahul Satija, James Nemesh, Karthik Shekhar, Melissa Goldman, Itay Tirosh, Allison R. Bialas, Nolan Kamitaki, Emily M. Martersteck, John J. Trombetta, David A. Weitz, Joshua R. Sanes, Alex K. Shalek, Aviv Regev, and Steven A. McCarroll. Highly parallel genome-wide expression profiling of individual cells using nanoliter droplets. Cell, 161:1202–1214, 5 2015.

[4] S. Steven Potter. Single-cell rna sequencing for the study of development, physiology and disease. Nature Reviews Nephrology, 14:479–492, 8 2018.

[5] Karthik A. Jagadeesh, Kushal K. Dey, Daniel T. Montoro, Rahul Mohan, Steven Gazal, Jesse M. Engreitz, Ramnik J. Xavier, Alkes L. Price, and Aviv Regev. Identifying disease-critical cell types and cellular processes by integrating single-cell rna-sequencing and human genetics. Nature Genetics, 54:1479–1492, 10 2022.

[6] David Lähnemann, Johannes Köster, Ewa Szczurek, Davis J. McCarthy, Stephanie C. Hicks, Mark D. Robinson, Catalina A. Vallejos, Kieran R. Campbell, Niko Beerenwinkel, Ahmed Mahfouz, Luca Pinello, Pavel Skums, Alexandros Stamatakis, Camille Stephan Otto Attolini, Samuel Aparicio, Jasmijn Baaijens, Marleen Balvert, Buys de Barbanson, Antonio Cappuccio, Giacomo Corleone, Bas E. Dutilh, Maria Florescu, Victor Guryev, Rens Holmer, Katharina Jahn, Thamar Jessurun Lobo, Emma M. Keizer, Indu Khatri, Szymon M. Kielbasa, Jan O. Korbel, Alexey M. Kozlov, Tzu Hao Kuo, Boudewijn P.F. Lelieveldt, Ion I. Mandoiu, John C. Marioni, Tobias Marschall, Felix Mölder, Amir Niknejad, Lukasz Raczkowski, Marcel Reinders, Jeroen de Ridder, Antoine Emmanuel Saliba, Antonios Somarakis, Oliver Stegle, Fabian J. Theis, Huan Yang, Alex Zelikovsky, Alice C. McHardy, Benjamin J. Raphael, Sohrab P. Shah, and Alexander Schönhuth. Eleven grand challenges in single-cell data science. Genome Biology, 21, 2 2020.

[7] Belinda Phipson, Luke Zappia, and Alicia Oshlack. Gene length and detection bias in single cell rna sequencing protocols. F1000Research, 6, 2017.

[8] Davide Risso, Katja Schwartz, Gavin Sherlock, and Sandrine Dudoit. Gc-content normalization for rna-seq data. BMC Bioinformatics, 12, 12 2011.

[9] Mark D Robinson, Davis J McCarthy, and Gordon K Smyth. edger: a bioconductor package for differential expression analysis of digital gene expression data. bioinformatics, 26(1):139–140, 2010.

[10] Peng Qiu. Embracing the dropouts in single-cell rna-seq analysis. Nature Communications, 11, 12 2020.

[11] Saket Choudhary and Rahul Satija. Comparison and evaluation of statistical error models for scrna-seq. Genome Biology, 23, 12 2022.

[12] Christoph Hafemeister and Rahul Satija. Normalization and variance stabilization of singlecell rna-seq data using regularized negative binomial regression. Genome Biology, 20, 12 2019.

[13] Wei Vivian Li and Jingyi Jessica Li. An accurate and robust imputation method scimpute for single-cell rna-seq data. Nature Communications, 9, 12 2018.

[14] Mengjie Chen and Xiang Zhou. Viper: Variability-preserving imputation for accurate gene expression recovery in single-cell rna sequencing studies. Genome Biology, 19, 11 2018.

[15] Mo Huang, Jingshu Wang, Eduardo Torre, Hannah Dueck, Sydney Shaffer, Roberto Bonasio, John I. Murray, Arjun Raj, Mingyao Li, and Nancy R. Zhang. Saver: Gene expression recovery for single-cell rna sequencing. Nature Methods, 15:539–542, 7 2018.

[16] George C Linderman, Jun Zhao, Manolis Roulis, Piotr Bielecki, Richard A Flavell, Boaz Nadler, and Yuval Kluger. Zero-preserving imputation of single-cell rna-seq data. Nature communications, 13(1):192, 2022.

[17] Gökcen Eraslan, Lukas M. Simon, Maria Mircea, Nikola S. Mueller, and Fabian J. Theis. Single-cell rna-seq denoising using a deep count autoencoder. Nature Communications, 10, 12 2019.

[18] Cédric Arisdakessian, Olivier Poirion, Breck Yunits, Xun Zhu, and Lana X. Garmire. Deep-impute: An accurate, fast, and scalable deep neural network method to impute single-cell rna-seq data. Genome Biology, 20, 10 2019.

[19] Efthymia Papalexi and Rahul Satija. Single-cell rna sequencing to explore immune cell heterogeneity. Nature Reviews Immunology, 18:35–45, 1 2018.

[20] Andre J. Faure, Jörn M. Schmiedel, and Ben Lehner. Systematic analysis of the determinants of gene expression noise in embryonic stem cells. Cell Systems, 5:471–484.e4, 11 2017.

[21] Xi Chen, Diederik P. Kingma, Tim Salimans, Yan Duan, Prafulla Dhariwal, John Schulman, Ilya Sutskever, and Pieter Abbeel. Variational lossy autoencoder. ArXiv, abs/1611.02731, 11 2016.

[22] Shengjia Zhao, Jiaming Song, and Stefano Ermon. Infovae: Information maximizing variational autoencoders. ArXiv, abs/1706.02262, 6 2017.

[23] Anders Boesen Lindbo Larsen, Søren Kaae Sønderby, Hugo Larochelle, and Ole Winther. Autoencoding beyond pixels using a learned similarity metric. ArXiv, abs/1512.09300, 12 2015.

[24] Andrew Butler, Paul Hoffman, Peter Smibert, Efthymia Papalexi, and Rahul Satija. Integrating single-cell transcriptomic data across different conditions, technologies, and species. Nature Biotechnology, 36:411–420, 6 2018.

[25] Matthijs J. Warrens and Hanneke Van Der Hoef. Understanding the Adjusted Rand Index and Other Partition Comparison Indices Based on Counting Object Pairs. Journal of Classification, 39(3):487–509, November 2022.

[26] Luis Aparicio, Mykola Bordyuh, Andrew J. Blumberg, and Raul Rabadan. A Random Matrix Theory Approach to Denoise Single-Cell Data. Patterns, 1(3):100035, June 2020.

[27] Iain M. Johnstone, Zongming Ma, Patrick O. Perry, and Morteza Shahram. RMTstat: Distributions, Statistics and Tests derived from Random Matrix Theory, 2022. R package version 0.3.1.

[28] Sara Aibar, Carmen Bravo Gonzalez-Blas, Thomas Moerman, Van Anh Huynh-Thu, Hana Imrichova, Gert Hulselmans, Florian Rambow, Jean-Christophe Marine, Pierre Geurts, Jan Aerts, Joost van den Oord, Zeynep Kalender Atak, Jasper Wouters, and Stein Aerts. Scenic: Single-cell regulatory network inference and clustering. Nature Methods, 14:1083– 1086, 2017.

[29] Tianzhi Wu, Erqiang Hu, Shuangbin Xu, Meijun Chen, Pingfan Guo, Zehan Dai, Tingze Feng, Lang Zhou, Wenli Tang, Li Zhan, Xiaocong Fu, Shanshan Liu, Xiaochen Bo, and Guangchuang Yu. clusterProfiler 4.0: A universal enrichment tool for interpreting omics data. The Innovation, 2(3):100141, August 2021.

[30] Francisco Avila Cobos, Mohammad Javad Najaf Panah, Jessica Epps, Xiaochen Long, Tsz-Kwong Man, Hua-Sheng Chiu, Elad Chomsky, Evgeny Kiner, Michael J Krueger, Diego di Bernardo, et al. Effective methods for bulk rna-seq deconvolution using scnrna-seq transcriptomes. Genome biology, 24(1):177, 2023.

[31] Yoav Benjamini and Yosef Hochberg. Controlling the false discovery rate: A practical and powerful approach to multiple testing. Journal of the Royal Statistical Society Series B: Statistical Methodology, 57:289–300, 1 1995.

[32] Yumei Li, Xinzhou Ge, Fanglue Peng, Wei Li, and Jingyi Jessica Li. Exaggerated false positives by popular differential expression methods when analyzing human population samples. Genome biology, 23(1):79, 2022.

[33] Amit Zeisel, Ana B. Muñoz-Manchado, Simone Codeluppi, Peter Lönnerberg, Gioele La Manno, Anna Juréus, Sueli Marques, Hermany Munguba, Liqun He, Christer Betsholtz, Charlotte Rolny, Gonçalo Castelo-Branco, Jens Hjerling-Leffler, and Sten Linnarsson. Cell types in the mouse cortex and hippocampus revealed by single-cell rna-seq. Science, 347:1138–1142, 3 2015.

[34] Grace X.Y. Zheng, Jessica M. Terry, Phillip Belgrader, Paul Ryvkin, Zachary W. Bent, Ryan Wilson, Solongo B. Ziraldo, Tobias D. Wheeler, Geoff P. McDermott, Junjie Zhu, Mark T. Gregory, Joe Shuga, Luz Montesclaros, Jason G. Underwood, Donald A. Masquelier, Stefanie Y. Nishimura, Michael Schnall-Levin, Paul W. Wyatt, Christopher M. Hindson, Rajiv Bharadwaj, Alexander Wong, Kevin D. Ness, Lan W. Beppu, H. Joachim Deeg, Christopher McFarland, Keith R. Loeb, William J. Valente, Nolan G. Ericson, Emily A. Stevens, Jerald P. Radich, Tarjei S. Mikkelsen, Benjamin J. Hindson, and Jason H. Bielas. Massively parallel digital transcriptional profiling of single cells. Nature Communications, 8, 1 2017.

[35] Xiaoping Han, Renying Wang, Yincong Zhou, Lijiang Fei, Huiyu Sun, Shujing Lai, Assieh Saadatpour, Ziming Zhou, Haide Chen, Fang Ye, et al. Mapping the mouse cell atlas by microwell-seq. Cell, 172(5):1091–1107, 2018.

[36] Florian Buettner, Kedar N. Natarajan, F. Paolo Casale, Valentina Proserpio, Antonio Scialdone, Fabian J. Theis, Sarah A. Teichmann, John C. Marioni, and Oliver Stegle. Computational analysis of cell-to-cell heterogeneity in single-cell rna-sequencing data reveals hidden subpopulations of cells. Nature Biotechnology, 33:155–160, 2015.

[37] Qinhuan Luo, Yaozhu Chen, and Xun Lan. Comse: analysis of single-cell rna-seq data using community detection-based feature selection. BMC Biology, 22, 12 2024.

[38] Antonio Scialdone, Kedar N. Natarajan, Luis R. Saraiva, Valentina Proserpio, Sarah A. Teichmann, Oliver Stegle, John C. Marioni, and Florian Buettner. Computational assignment of cell-cycle stage from single-cell transcriptome data. Methods, 85:54–61, 9 2015.

[39] Itay Tirosh, Benjamin Izar, Sanjay M. Prakadan, Marc H. Wadsworth, Daniel Treacy, John J. Trombetta, Asaf Rotem, Christopher Rodman, Christine Lian, George Murphy, Mohammad Fallahi-Sichani, Ken Dutton-Regester, Jia-Ren Lin, Ofir Cohen, Parin Shah, Diana Lu, Alex S. Genshaft, Travis K. Hughes, Carly G. K. Ziegler, Samuel W. Kazer, Aleth Gaillard, Kellie E. Kolb, Alexandra-Chloé Villani Cory M. Johannessen, Aleksandr Y. Andreev, Eliezer M. Van Allen, Monica Bertagnolli, Peter K. Sorger, Ryan J. Sullivan, Keith T. Flaherty, Dennie T. Frederick, Judit Jané-Valbuena, Charles H. Yoon, Orit Rozenblatt-Rosen, Alex K. Shalek, Aviv Regev, and Levi A. Garraway. Dissecting the multicellular ecosystem of metastatic melanoma by single-cell rna-seq. Science, 352:189– 196, 4 2016.

[40] Aaron T.L. Lun, Karsten Bach, and John C. Marioni. Pooling across cells to normalize single-cell rna sequencing data with many zero counts. Genome Biology, 17, 4 2016.

[41] Tallulah S. Andrews and Martin Hemberg. M3drop: Dropout-based feature selection for scrnaseq. Bioinformatics, 35:2865–2867, 8 2019.

[42] Malte D. Luecken, M. Büttner, K. Chaichoompu, A. Danese, M. Interlandi, M. F. Mueller, D. C. Strobl, L. Zappia, M. Dugas, M. Colomé-Tatché, and Fabian J. Theis. Benchmarking atlas-level data integration in single-cell genomics. Nature Methods, 19:41–50, 1 2022.

[43] Ilya Korsunsky, Nghia Millard, Jean Fan, Kamil Slowikowski, Fan Zhang, Kevin Wei, Yuriy Baglaenko, Michael Brenner, Po ru Loh, and Soumya Raychaudhuri. Fast, sensitive and accurate integration of single-cell data with harmony. Nature Methods, 16:1289–1296, 12 2019.

[44] Michael I Love, Wolfgang Huber, and Simon Anders. Moderated estimation of fold change and dispersion for rna-seq data with deseq2. Genome biology, 15(12):1–21, 2014.

[45] Matthew E Ritchie, Belinda Phipson, D. Wu, Yifang Hu, Charity W Law, Wei Shi, and Gordon K Smyth. limma powers differential expression analyses for rna-sequencing and microarray studies. Nucleic acids research, 43(7):e47, 2015.

[46] Luba Nalysnyk, Javier Cid-Ruzafa, Philip Rotella, and Dirk Esser. Incidence and prevalence of idiopathic pulmonary fibrosis: Review of the literature. European Respiratory Review, 21:355–361, 12 2012.

[47] Vishwaraj Sontake, Rajesh K. Kasam, Debora Sinner, Thomas R. Korfhagen, Geereddy B. Reddy, Eric S. White, Anil G. Jegga, and Satish K. Madala. Wilms’ tumor 1 drives fibroproliferation and myofibroblast transformation in severe fibrotic lung disease. JCI insight, 3, 8 2018.

[48] Michael J. Bollong, Baiyuan Yang, Naja Vergani, Brittney A. Beyer, Emily N. Chin, Claudio Zambaldo, Danling Wang, Arnab K. Chatterjee, Luke L. Lairson, and Peter G. Schultz. Small molecule-mediated inhibition of myofibroblast transdifferentiation for the treatment of fibrosis. Proceedings of the National Academy of Sciences of the United States of America, 114:4679–4684, 5 2017.

[49] Gil Stelzer, Naomi Rosen, Inbar Plaschkes, Shahar Zimmerman, Michal Twik, Simon Fishilevich, Tsippi Iny Stein, Ron Nudel, Iris Lieder, Yaron Mazor, Sergey Kaplan, Dvir Dahary, David Warshawsky, Yaron Guan-Golan, Asher Kohn, Noa Rappaport, Marilyn Safran, and Doron Lancet. The genecards suite: From gene data mining to disease genome sequence analyses. Current Protocols in Bioinformatics, 2016:1.30.1–1.30.33, 2016.

[50] Tian Tian, Ji Wan, Qi Song, and Zhi Wei. Clustering single-cell rna-seq data with a model-based deep learning approach. Nature Machine Intelligence, 1:191–198, 4 2019.

[51] Yang Li, Mingcong Wu, Shuangge Ma, and Mengyun Wu. Zinbmm: a general mixture model for simultaneous clustering and gene selection using single-cell transcriptomic data. Genome Biology, 24, 12 2023.

[52] Yanyan Zeng, Jing Li, Chaochun Wei, Hongyu Zhao, and Tao Wang. mbdenoise: microbiome data denoising using zero-inflated probabilistic principal components analysis. Genome Biology, 23(1):94, 2022.

[53] Hantao Shu, Jingtian Zhou, Qiuyu Lian, Han Li, Dan Zhao, Jianyang Zeng, and Jianzhu Ma. Modeling gene regulatory networks using neural network architectures. Nature Computational Science, 1:491–501, 7 2021.

[54] Dominic Grün, Lennart Kester, and Alexander Van Oudenaarden. Validation of noise models for single-cell transcriptomics. Nature Methods, 11:637–640, 2014.

[55] Fan Guo, Lin Li, Jingyun Li, Xinglong Wu, Boqiang Hu, Ping Zhu, Lu Wen, and Fuchou Tang. Single-cell multi-omics sequencing of mouse early embryos and embryonic stem cells. Cell Research, 27:967–988, 8 2017.

