## Supplementary Materials for "Denoising Single-Cell RNA-Seq Data with a Deep Learning-Embedded Statistical Framework"

**2 Additional Figures 9**

### 1 Supplementary Material

#### 1.1 Details on infoVAE-GAN model

To select the hyperparameters $\gamma_1$ and $\gamma_2$, we explored various configurations with $\gamma_1 \in \{1, 2, 5\}$ and $\gamma_2 \in \{0.1, 0.5, 1\}$. We observed consistent group structures $U$ and $V$ across these different settings, highlighting the robust performance of the infoVAE-GAN model across diverse datasets. Based on these experiments, we finalized our choice to $\gamma_1 = 2$ and $\gamma_2 = 0.5$.

Furthermore, to improve the model's representational capacity and generalization, we employed an ensemble approach involving multiple identical encoders. The outputs from these encoders were averaged to yield a robust latent representation $x$. We empirically assessed ensembles of size between 10 and 50 encoders, and found that the performance of infoVAE-GAN remained stable, demonstrating insensitivity to the specific ensemble size.

#### 1.2 Details on EM algorithms

We aim to maximize the following complete log-likelihood, given a fixed latent factor matrix $V$:

$$\begin{aligned}\mathcal{L}(\Delta \mid \Omega, Z) = &\sum_{ij}(z_{ij}\log\phi_j + (1-z_{ij})\log((1-\phi_j)\,\mathbb{P}(y_{ij} \mid \mu_{ij}, \theta_j)))\\ &-\lambda_u \sum_j u_j^\top u_j - \lambda_\zeta \sum_j \zeta_j^2 - \lambda_\xi \sum_i \xi_i^2,\end{aligned}$$

where $\Omega = \{Y,\ V\}$, $\Delta = \{\phi_j,\ \theta_j,\ \xi_i,\ \zeta_j,\ \alpha_j,\ \beta_i,\ u_j \mid i \in \{1, \cdots, N\},\ j \in \{1, \cdots, M\}\}$, $y_{ij}$ are the observed counts, $z_{ij}$ are latent indicators for dropout events, and $\lambda_u, \lambda_\zeta, \lambda_\xi$ are regularization hyperparameters. The mean parameter is expressed as $\log\mu_{ij} = \xi_i + \zeta_j + v_i^\top\alpha_j + \beta_i^\top u_j$.

The parameters $\Delta$ are estimated using an EM algorithm. In the E-step, the posterior probabilities of the latent variables $Z$ at iteration $t$, $q^{(t)}(Z) = p(Z \mid \Omega, \Delta^{(t-1)})$, are computed as

$$\begin{aligned}&\mathbb{E}_{q^{(t)}}[z_{ij} \mid y_{ij}, V, \Delta^{(t-1)}]\\ =\ &\mathbb{P}(z_{ij} = 1 \mid y_{ij}, V, \Delta^{(t-1)})\\ =\ &\frac{\mathbb{P}(y_{ij} \mid z_{ij} = 1, V, \Delta^{(t-1)})\,\mathbb{P}(z_{ij} = 1 \mid V, \Delta^{(t-1)})}{\mathbb{P}(y_{ij} \mid z_{ij} = 1, V, \Delta^{(t-1)})\,\mathbb{P}(z_{ij} = 1 \mid V, \Delta^{(t-1)}) + \mathbb{P}(y_{ij} \mid z_{ij} = 0, V,\ \Delta^{(t-1)})\,\mathbb{P}(z_{ij} = 0 \mid V, \Delta^{(t-1)})}\\ =\ &\frac{\mathrm{I}\{y_{ij} = 0\}\phi_j^{(t-1)}}{\mathrm{I}\{y_{ij} = 0\}\phi_j^{(t-1)} + \mathbb{P}(y_{ij} \mid V, \Delta^{(t-1)})(1-\phi_j^{(t-1)})},\end{aligned}$$

or equivalently,

$$z_{ij}^{(t)} = \frac{\mathrm{I}\{y_{ij} = 0\}\phi_j^{(t-1)}}{\mathrm{I}\{y_{ij} = 0\}\phi_j^{(t-1)} + \mathbb{P}(y_{ij} \mid \Delta^{(t-1)})(1 - \phi_j^{(t-1)})}.$$

The corresponding Q-function (or Evidence Lower Bound, ELBO) is:

$$\begin{aligned}
Q^{(t)}(\Delta \mid \Omega, Z) &= \mathbb{E}_{q^{(t)}}[\mathcal{L}(\Delta \mid \Omega, Z)] \\
&= \sum_{ij} \left( z_{ij}^{(t)} \log \phi_j + \left(1 - z_{ij}^{(t)}\right) \log((1 - \phi_j)\, \mathbb{P}(y_{ij} \mid \mu_{ij}, \theta_j)) \right) \\
&\quad - \lambda_u \sum_j u_j^T u_j - \lambda_\zeta \sum_j \zeta_j^2 - \lambda_\xi \sum_i \xi_i^2 \\
&= \sum_{ij} \left( z_{ij}^{(t)} \log \phi_j + \left(1 - z_{ij}^{(t)}\right) \log(1 - \phi_j) \right) \\
&\quad + \sum_{ij} \left(1 - z_{ij}^{(t)}\right) \log \left( \frac{\Gamma\left(y_{ij} + \frac{1}{\theta_j}\right)}{\Gamma(y_{ij} + 1)\Gamma\left(\frac{1}{\theta_j}\right)} (1 + \theta_j \mu_{ij})^{-\frac{1}{\theta_j}} \left( \frac{\theta_j \mu_{ij}}{1 + \theta_j \mu_{ij}} \right)^{y_{ij}} \right) \\
&\quad - \lambda_u \sum_j u_j^T u_j - \lambda_\zeta \sum_j \zeta_j^2 - \lambda_\xi \sum_i \xi_i^2.
\end{aligned}$$

In the M-step, parameters are updated to maximize the above Q-function:

$$\Delta^{(t)} = \underset{\Delta}{\operatorname{argmax}}\, Q^{(t)}(\Delta \mid \Omega, Z).$$

The parameters are updated as follows. We first update $\phi_j$ by

$$\partial_{\phi_j} Q^{(t)} = \sum_i \frac{z_{ij}^{(t)}}{\phi_j} - \frac{1 - z_{ij}^{(t)}}{1 - \phi_j} = 0 \;\Rightarrow \phi_j^{(t)} = \frac{1}{N} \sum_i z_{ij}^{(t)},$$

and $\theta_j$ by solving the following equation with all the other parameters fixed

$$\partial_{\theta_j} Q^{(t)} = \frac{1}{\theta_j^2} \sum_i \left(1 - z_{ij}^{(t)}\right) \left[ \psi\left(\frac{1}{\theta_j}\right) - \psi\left(y_{ij} + \frac{1}{\theta_j}\right) + \log(1 + \theta_j \mu_{ij}) + \frac{\theta_j (y_{ij} - \mu_{ij})}{1 + \theta_j \mu_{ij}} \right] = 0,$$

where $\psi(\cdot)$ is digamma function. Then we update $\xi_i,\ \zeta_j,\ \alpha_j,\ \beta_i$ according to the following score functions

$$
\begin{aligned}
\partial_{\xi_i} Q^{(t)} &= \sum_j \partial_{\mu_{ij}} Q^{(t)} \mu_{ij} - 2\lambda_\xi \xi_i, \\
\partial_{\zeta_j} Q^{(t)} &= \sum_i \partial_{\mu_{ij}} Q^{(t)} \mu_{ij} - 2\lambda_\zeta \zeta_j, \\
\partial_{\alpha_j} Q^{(t)} &= \sum_i \partial_{\mu_{ij}} Q^{(t)} \mu_{ij} v_i, \\
\partial_{\beta_i} Q^{(t)} &= \sum_j \partial_{\mu_{ij}} Q^{(t)} \mu_{ij} u_j,
\end{aligned}
$$

where

$$
\partial_{\mu_{ij}} Q^{(t)} = \left(1 - z_{ij}^{(t)}\right) \frac{y_{ij} - \mu_{ij}}{\mu_{ij}\left(1 + \theta_j \mu_{ij}\right)} \Rightarrow \partial_{\mu_{ij}} Q^{(t)} \mu_{ij} = \left(1 - z_{ij}^{(t)}\right) \frac{y_{ij} - \mu_{ij}}{1 + \theta_j \mu_{ij}}.
$$

Finally, we shall update the estimation of latent factors with the score function as

$$
\partial_{u_j} Q^{(t)} = \sum_i \partial_{\mu_{ij}} Q^{(t)} \mu_{ij} \beta_i - 2\lambda_u u_j.
$$

These steps are iterated until convergence to obtain the final estimators $\hat{\Delta}$. Practically, the latent factor matrix $U$ can be fixed to expedite computation by omitting its update during iterations.

Let the final estimators be $\hat{\Delta} = \{\hat{\phi}_j,\ \hat{\theta}_j,\ \hat{\xi}_i,\ \hat{\zeta}_j,\ \hat{\alpha}_j,\ \hat{\beta}_i,\ \hat{u}_j \mid i \in \{1, \cdots, N\},\ \ j \in \{1, \cdots, M\}\}$. We have the primary estimation of mean parameter $\hat{\mu}_{ij}$ as

$$
\log \hat{\mu}_{ij} = \hat{\xi}_i + \hat{\zeta}_j + v_i^\top \hat{\alpha}_j + \hat{\beta}_i^\top \hat{u}_j,
$$

and the denoised estimation by dropping $\hat{\xi}_i + \hat{\zeta}_j$ as

$$
\log \hat{\mu}_{ij}^* = v_i^\top \hat{\alpha}_j + \hat{\beta}_i^\top \hat{u}_j.
$$

##### 1.3 Confusion matrix and external clustering evaluation

We first present the widely-used confusion matrix in Table 1, which summarizes the agreement between predicted and true binary labels through four outcomes: true positives (TP), false negatives (FN), false positives (FP), and true negatives (TN). These outcomes form the

| | | Prediction | |
|---|---|---|---|
| | | 1 | 0 |
| True | 1 | TP | FN |
| | 0 | FP | TN |

Table 1: Confusion matrix (TP = true positive, FN = false negative, FP = false positive, TN = true negative).

basis for various standard evaluation metrics used to assess predictive accuracy in differential expression genes (DEGs) analysis as described in Section 3.3.

1. Precision:

$$Precision = \frac{TP}{TP + FP};$$

2. False Discovery Rate (FDR)

$$FDR = \frac{FP}{TP + FP} = 1 - Precision;$$

3. Specificity

$$Specificity = \frac{TN}{TN + FP};$$

4. False Postive Rate (FPR)

$$FPR = \frac{FP}{FP + TN} = 1 - Specificity;$$

5. Accuracy (ACC)

$$ACC = \frac{TP + TN}{TP + FN + FP + TN}.$$

Using FPR and True Positive Rate (TPR), we construct the Receiver Operating Characteristic (ROC) curve. The area under this curve (AUC) is a widely recognized metric reflecting overall classifier performance.

We further detail the metrics used for external clustering evaluation in Section 3.1. Consider a dataset with $N$ samples, where two distinct partitions exist: $P = P_1, \ldots, P_K$ for true labels and $Q = Q_1, \ldots, Q_L$ for predicted labels. Let $n_{ij} = |P_i \cap Q_j|$ represent the number of samples common to clusters $P_i$ and $Q_j$. Define $n_{i\cdot} = \sum_{j=1}^{L} n_{ij}$ and $n_{\cdot j} = \sum_{i=1}^{K} n_{ij}$ as the total samples in clusters $P_i$ and $Q_j$, respectively. The confusion matrix entries for pairs of samples are defined as follows:

$$
\begin{aligned}
TP &= \frac{1}{2}\left(\sum_{i=1}^{K}\sum_{j=1}^{L} n_{ij}^2 - N\right);\\
FN &= \frac{1}{2}\left(\sum_{i=1}^{K} n_{i\cdot}^2 - \sum_{i=1}^{K}\sum_{j=1}^{L} n_{ij}^2\right);\\
FP &= \frac{1}{2}\left(\sum_{j=1}^{L} n_{\cdot j}^2 - \sum_{i=1}^{K}\sum_{j=1}^{L} n_{ij}^2\right);\\
TN &= \frac{1}{2}\left(N^2 + \sum_{i=1}^{K}\sum_{j=1}^{L} n_{ij}^2 - \sum_{i=1}^{K} n_{i\cdot}^2 - \sum_{j=1}^{L} n_{\cdot j}^2\right);\\
Total &= TP + FN + FP + TN.
\end{aligned}
$$

Using the formulation above, we have

1. Purity:

$$Purity = \frac{1}{N}\sum_{j=1}^{L} \max_{j} n_{ij};$$

2. F score (or Dice index): let $Recall = \frac{TP}{TP+FN}$ as above, then

$$F\ score = \frac{2 * Precision * Racall}{Precision + Recall} = \frac{2TP}{2TP + FP + FN};$$

3. Rand Index (RI):

$$RI = \frac{TP + TN}{Total};$$

4. Adjusted Rand Index (ARI):

$$ARI = \frac{2(TP * TN - FP * FN)}{(TP + FN)(FN + TN) + (TP + FP)(FP + TN)}.$$

For evaluating sets characterized by additional metrics (FDR, logFC, $f$ values, and so on), we utilize classic statistical metrics such as Pearson correlation, Wasserstein-2 distance, and cross entropy. For instance, comparing $f$ values in true DEGs (tDEGs) and predicted DEGs (pDEGs), we calculate

$$Corr_f = \frac{\sum_{g\in G}(f_g - \bar{f})(\hat{f}_g - \bar{\hat{f}})}{\sqrt{\sum_{g\in G}(f_g - \bar{f})^2}\sqrt{\sum_{g\in G}(\hat{f}_g - \bar{\hat{f}})^2}},$$

where $G$ is the intersection between tDEG and pDEG sets, $f_g$ is the $f$ value in tDEG set and $\hat{f}_g$ is the $f$ value in pDEG set. $\bar{f}$ and $\bar{\hat{f}}$ is their mean values respectively.

$$Wass_f = \sqrt{\int_0^1 \left|F_{G,f}^{-1}(t) - F_{G,\hat{f}}^{-1}(t)\right|^2 dt},$$

where $F_{G,f}^{-1}(t) = \inf_x\{F_{G,f}(x) \geqslant t\}$ is empirical quantile functions and $F_{G,f}(x) = \frac{1}{|G|}\sum_{i=1}^{n} I\{f_g \leqslant t\}$ is the empirical cumulative distribution function of the $f$ value in tDEG set. Similar formula $F_{G,\hat{f}}^{-1}(t)$ for $f$ value in pDEG set. For cross-entropy evaluation, the entire gene set is utilized, assigning tDEGs a value of 1 and non-tDEGs a value of 0. The cross-entropy is calculated as:

$$CE_f = -\sum_{g\in G}[f_g \log(\hat{f}_g) + (1 - f_g)\log(1 - \hat{f}_g)],$$

where $G$ denotes all genes in a bootstrap simulation.

##### 1.4 Random matrix statistics

Consider a matrix $X = (x_1, \cdots, x_n) \in \mathbb{R}^{D\times N}$, where each entry of $X$ is independently sampled from a Gaussian distribution $N(0, \sigma^2)$. The sample covariance matrix (i.e., Wishart matrix) is defined as

$$W = \frac{1}{N}XX^\top = \frac{1}{N}\sum_{i=1}^{N} x_i x_i^\top.$$

The following eigenvalue-based statistics are primarily utilized:

1. **Empirical spectral density**: Defined as the empirical spectral measure:

$$\mu_W = \frac{1}{D}\sum_{i=1}^{D}\delta_{\lambda_i}, \tag{1}$$

where $\lambda_i$ are the eigenvalues of $W$, and $\delta_x(t) = \mathrm{I}\{t \geqslant x\}$. Then the empirical spectral measure converges to the so-called Marchenko-Pastur distribution when $N \to +\infty$, $D \to$

$+\infty$, $\gamma = D/N \in (0, 1]$ with the following density

$$\rho_{MP}(t) = \frac{1}{2\pi\gamma\sigma^2}\frac{\sqrt{(\lambda_+ - t)(t - \lambda_-)}}{t}\mathrm{I}[\lambda_-, \lambda_+],\ \lambda_\pm = \sigma^2(1 \pm \sqrt{\gamma})^2. \tag{2}$$

2. **Empirical normalized level-spacing density**: Consider eigenvalues ordered as $\lambda_{(D)} \geqslant \ldots \geqslant \lambda_{(1)}$ and define the gaps $d_i = \lambda_{(i+1)} - \lambda_{(i)}$ for $i \in \{1, \ldots, D-1\}$. The normalized gaps are $s_i = \frac{d_i}{\bar{d}}$, where $\bar{d} = (D-1)^{-1}\sum_{i=1}^{D-1} d_i$ is the mean gap. The empirical normalized level-spacing measure is defined as:

$$\mu_G = \frac{1}{D-1}\sum_{i=1}^{D-1}\delta_{s_i}. \tag{3}$$

Under the same asymptotic conditions as above, this distribution approaches the Wigner Surmise:

$$\rho_{WS}(t) = \frac{\pi t}{2}e^{-\frac{\pi}{4}t^2}\mathrm{I}[0, +\infty). \tag{4}$$

3. **Spectral radius**: The distribution of the largest eigenvalue, known as the spectral radius of the Wishart matrix, follows the Tracy-Widom distribution asymptotically:

$$\mathbb{P}(\lambda_{(D)} \leqslant t) = F_1(D^{\frac{2}{3}}(\lambda_{(D)} - t)), \tag{5}$$

where $F_1(\cdot)$ is the Tracy-Widom distribution corresponding to $\beta = 1$. Specifically, it is given by:

$$F_1(t) = \exp\left\{-\frac{1}{2}\int_t^\infty \big(q(x) + (x-t)q^2(x)\big)\,dx\right\},$$

with $q(x)$ satisfying the Painlevé II differential equation:

$$q''(x) = xq(x) + 2q(x)^3,$$

and boundary condition $q(x) \sim \mathrm{Ai}(x)$ as $x \to +\infty$, where $\mathrm{Ai}(x)$ is the Airy function:

$$\mathrm{Ai}(x) = \frac{1}{\pi}\int_0^\infty \cos\left(\frac{s^3}{3} + xs\right)ds.$$

#### 2 Additional Figures

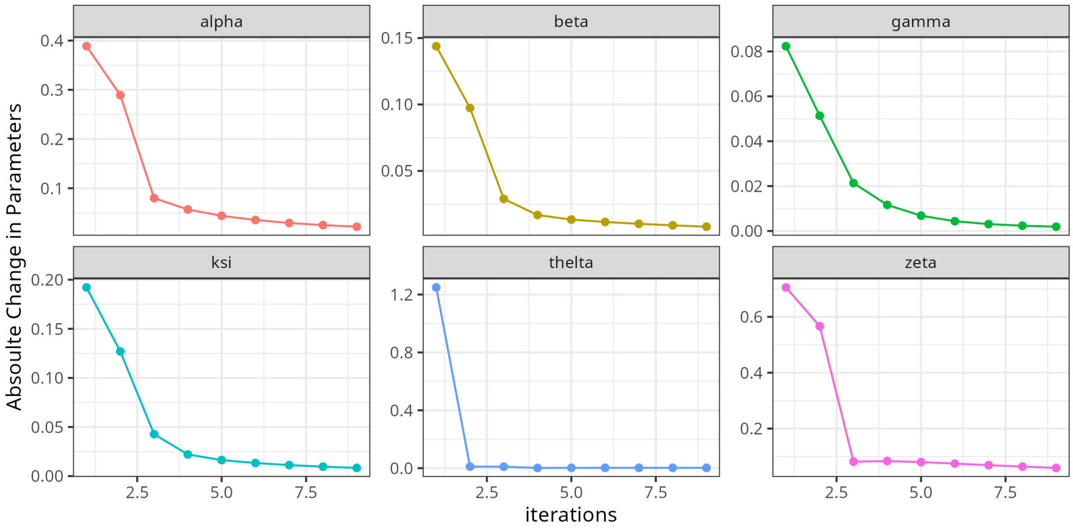

Figure S1: Convergence of parameters Δ estimated by the EM algorithm.

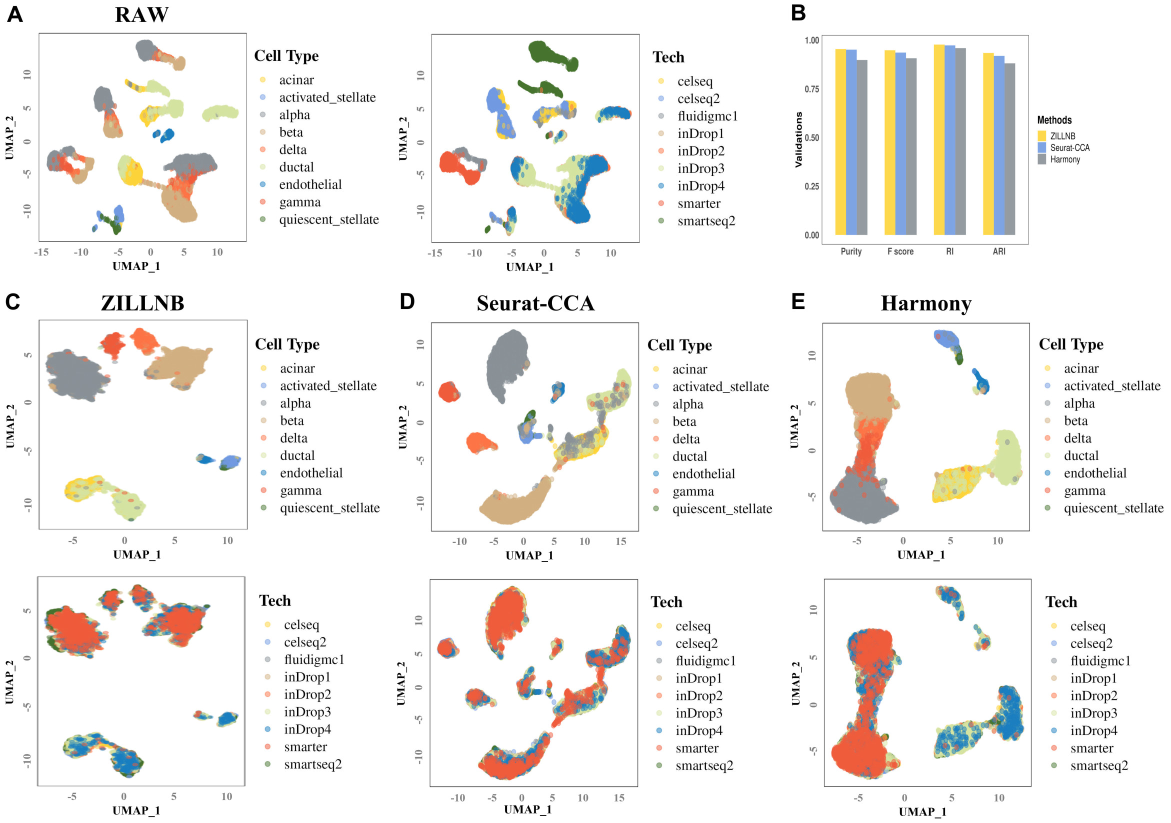

Figure S2: Latent cell factor matrix $V$ of ZILLNB achieves considerable performance on batch-effect removal compared to other commonly used methods (Seurat-CCA and Harmony) illustrated with human pancreatic islet datasets. All plots are labeled by cell types and scRNA-seq protocols, including CEL-Seq [3, 4], CEL-Seq2 [7, 2], Fluidigm C1 [6, 9], InDrop 1-4 [1, 5], Smart-Seq [11, 10], and Smart-Seq2 [12, 8]. (A) UMAP plots of raw human pancreatic islet datasets without batch removal. Data preprocessing involved extracting the top 2,000 highly variable genes, followed by dimension reduction using the first 10 principal components with UMAP. We can observe greater variability between different scRNA-seq protocols relative to cell types. (B) Clustering validation metrics (Purity, F score, RI, ARI) comparing performance and robustness of ZILLNB against Seurat-CCA and Harmony methods. (C, D, E) UMAP plots of integrated human pancreatic scRNA-seq data utilizing the biological component of the latent factor matrix derived from the ZILLNB model, Seurat-CCA, and Harmony. The last two methods use the top 2,000 genes selected by Seurat.

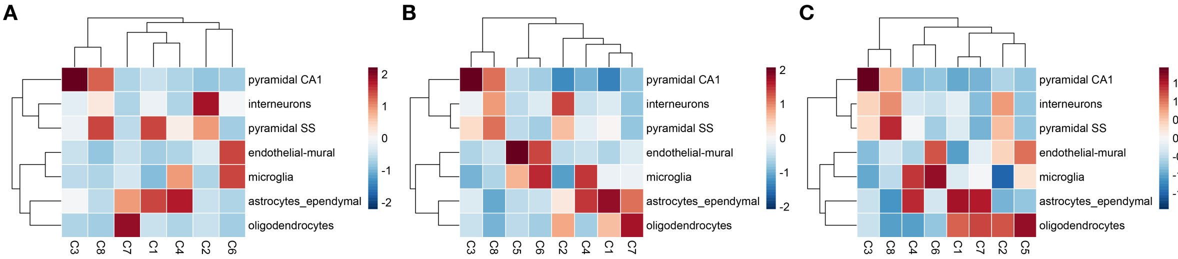

Figure S3: Heatmap illustrating the enrichment of marker genes between predicted gene clusters and cell types. Panels (A), (B), and (C) present results using the top 100, 500, and all marker genes selected by the Seurat method, respectively.

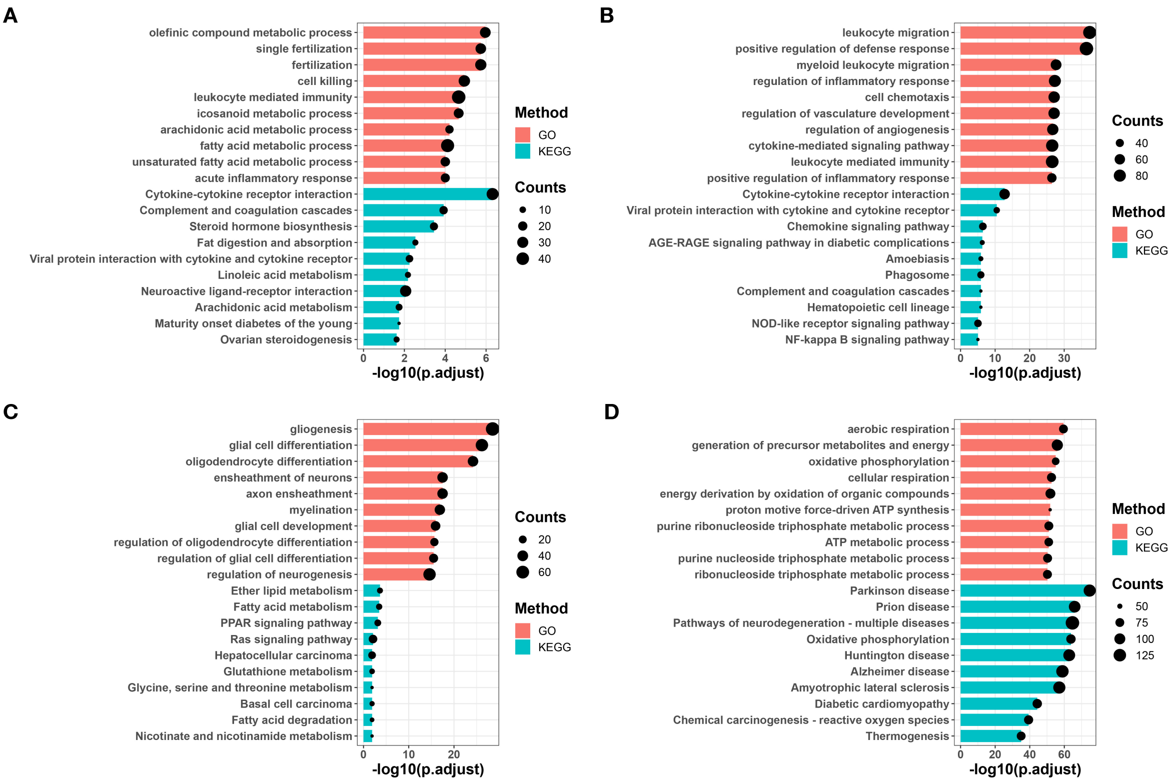

Figure S4: Gene set enrichment analysis results using GO and KEGG databases across different gene clusters. Panels (A), (B), (C), and (D) correspond to clusters 5, 6, 7, and 8, respectively. Box plots represent the negative logarithm of adjusted $p$-values, while the size of black points indicates the number of genes intersecting each specific GO or KEGG term with the respective gene cluster.

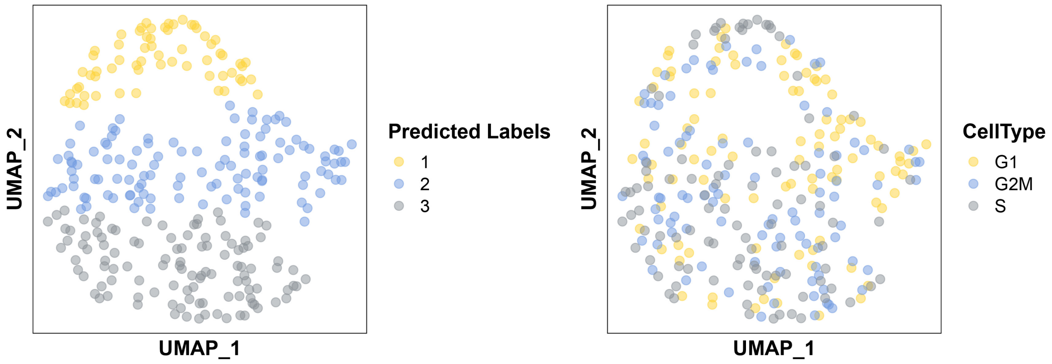

Figure S5: UMAP visualization of the HSC cell cycle dataset using the ZILLNB method's $\exp(\hat{\alpha}^\top V)$ matrix with predicted labels and true cell types.

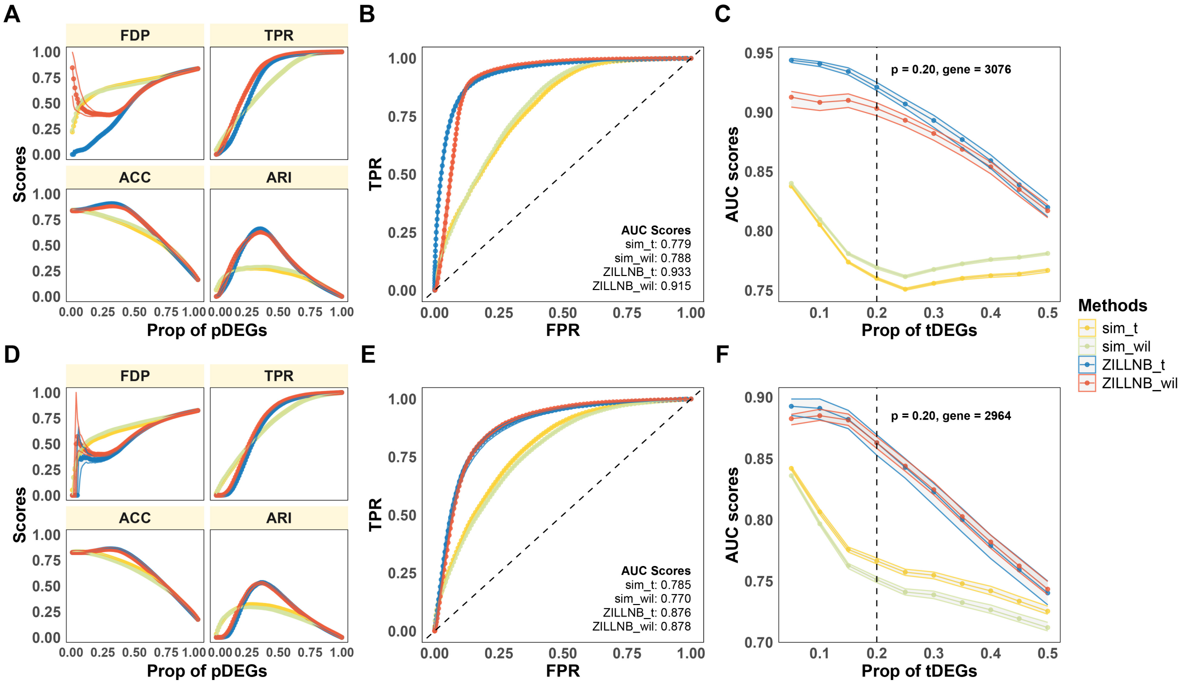

Figure S6: ROC curves and AUC scores from bootstrap experiments evaluating DEGs selection using tumor cell line T47D versus non-tumor cell lines Jurkat (first row) and Thp1 (second row). (A, D) FDP, TPR, ACC, and ARI metrics; (B, E) ROC curves with fixed true DEGs (tDEGs) proportions; (C, F) AUC scores across varying tDEGs proportions (0–0.5). Dashed lines represent scenarios with approximately 3,000 tDEGs (top 20%). "sim_t/sim_wil" uses Seurat's log-normalized data with t/Wilcoxon tests; "ZILLNB_t/ZILLNB_wil" uses the denoised ZILLNB matrix with t/Wilcoxon tests.

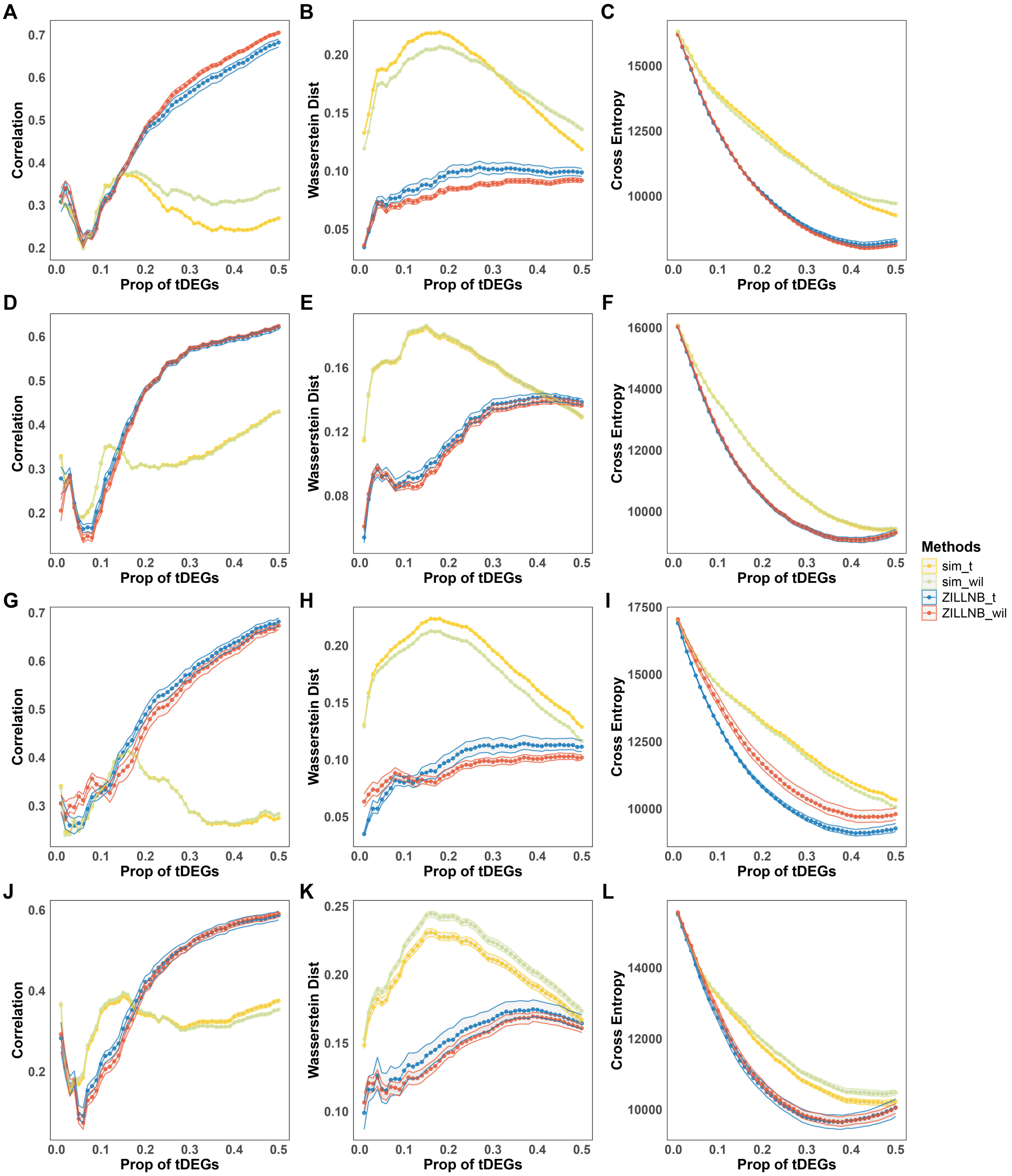

Figure S7: Similarity metrics (Pearson correlation, Wasserstein-2 distance, cross entropy) between bulk RNA-seq and scRNA-seq datasets from bootstrap DEG selection experiments. Rows represent different tumor/non-tumor comparisons (BT474/Jurkat, BT474/Thp1, T47D/Jurkat, T47D/Thp1). Methods compared include "sim_t/sim_wil" (Seurat log-normalized data with t/Wilcoxon tests) and "ZILLNB_t/ZILLNB_wil" (ZILLNB-denoised data with t/Wilcoxon tests).

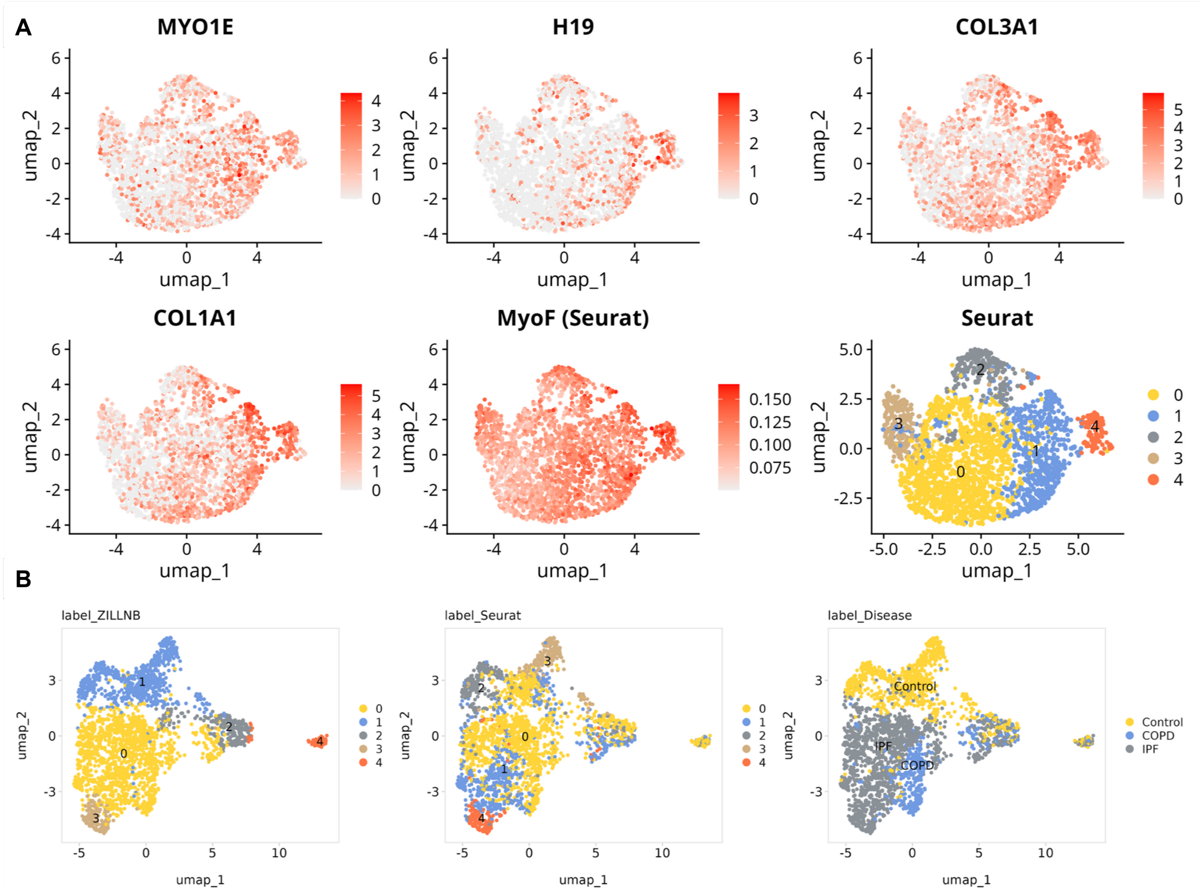

Figure S8: (A) UMAP plots of fibroblasts from IPF, COPD, and healthy controls, displaying myofibroblast marker expression, AUCell scores, and Seurat-predicted labels. (B) UMAP plots with predicted labels from ZILLNB-denoised data, annotated by disease type.

#### References

[1] Maayan Baron, Adrian Veres, Samuel L. Wolock, Aubrey L. Faust, Renaud Gaujoux, Amedeo Vetere, Jennifer Hyoje Ryu, Bridget K. Wagner, Shai S. Shen-Orr, Allon M. Klein, Douglas A. Melton, and Itai Yanai. A single-cell transcriptomic map of the human and mouse pancreas reveals inter- and intra-cell population structure. Cell Systems, 3:346–360.e4, 10 2016.

[2] Dominic Grün, Lennart Kester, and Alexander Van Oudenaarden. Validation of noise models for single-cell transcriptomics. Nature Methods, 11:637–640, 2014.

[3] Dominic Grün, Mauro J. Muraro, Jean Charles Boisset, Kay Wiebrands, Anna Lyubimova, Gitanjali Dharmadhikari, Maaike van den Born, Johan van Es, Erik Jansen, Hans Clevers, Eelco J.P. de Koning, and Alexander van Oudenaarden. De novo prediction of stem cell identity using single-cell transcriptome data. Cell Stem Cell, 19:266–277, 8 2016.

[4] Tamar Hashimshony, Florian Wagner, Noa Sher, and Itai Yanai. Cel-seq: Single-cell rna-seq by multiplexed linear amplification. Cell Reports, 2:666–673, 9 2012.

[5] Allon M. Klein, Linas Mazutis, Ilke Akartuna, Naren Tallapragada, Adrian Veres, Victor Li, Leonid Peshkin, David A. Weitz, and Marc W. Kirschner. Droplet barcoding for single-cell transcriptomics applied to embryonic stem cells. Cell, 161:1187–1201, 5 2015.

[6] Nathan Lawlor, Joshy George, Mohan Bolisetty, Romy Kursawe, Lili Sun, V. Sivakamasundari, Ina Kycia, Paul Robson, and Michael L. Stitzel. Single-cell transcriptomes identify human islet cell signatures and reveal cell-type-specific expression changes in type 2 diabetes. Genome Research, 27:208–222, 2 2017.

[7] Mauro J. Muraro, Gitanjali Dharmadhikari, Dominic Grün, Nathalie Groen, Tim Dielen, Erik Jansen, Leon van Gurp, Marten A. Engelse, Francoise Carlotti, Eelco J.P. de Koning, and Alexander van Oudenaarden. A single-cell transcriptome atlas of the human pancreas. Cell Systems, 3:385–394.e3, 10 2016.

[8] Simone Picelli, Åsa K. Björklund, Omid R. Faridani, Sven Sagasser, Gösta Winberg, and Rickard Sandberg. Smart-seq2 for sensitive full-length transcriptome profiling in single cells. Nature Methods, 10:1096–1100, 11 2013.

[9] Alex A. Pollen, Tomasz J. Nowakowski, Joe Shuga, Xiaohui Wang, Anne A. Leyrat, Jan H. Lui, Nianzhen Li, Lukasz Szpankowski, Brian Fowler, Peilin Chen, Naveen Ramalingam, Gang Sun, Myo Thu, Michael Norris, Ronald Lebofsky, Dominique Toppani, Darnell W. Kemp, Michael Wong, Barry Clerkson, Brittnee N. Jones, Shiquan Wu, Lawrence Knutsson, Beatriz Alvarado, Jing Wang, Lesley S. Weaver, Andrew P. May, Robert C. Jones, Marc A. Unger, Arnold R. Kriegstein, and Jay A.A. West. Low-coverage single-cell mrna sequencing reveals cellular heterogeneity and activated signaling pathways in developing cerebral cortex. Nature Biotechnology, 32:1053–1058, 10 2014.

[10] Daniel Ramsköld, Shujun Luo, Yu Chieh Wang, Robin Li, Qiaolin Deng, Omid R. Faridani, Gregory A. Daniels, Irina Khrebtukova, Jeanne F. Loring, Louise C. Laurent, Gary P. Schroth, and Rickard Sandberg. Full-length mrna-seq from single-cell levels of rna and individual circulating tumor cells. Nature Biotechnology, 30:777–782, 2012.

[11] Yurong Xin, Jinrang Kim, Haruka Okamoto, Min Ni, Yi Wei, Christina Adler, Andrew J. Murphy, George D. Yancopoulos, Calvin Lin, and Jesper Gromada. Rna sequencing of single human islet cells reveals type 2 diabetes genes. Cell Metabolism, 24:608–615, 10 2016.

[12] Åsa Segerstolpe, Athanasia Palasantza, Pernilla Eliasson, Eva Marie Andersson, Anne Christine Andréasson, Xiaoyan Sun, Simone Picelli, Alan Sabirsh, Maryam Clausen, Magnus K. Bjursell, David M. Smith, Maria Kasper, Carina Ämmälä, and Rickard Sandberg. Single-cell transcriptome profiling of human pancreatic islets in health and type 2 diabetes. Cell Metabolism, 24:593–607, 10 2016.
